# Parallel Mosaic Speciation via Mutation-order and Ecological Divergence

**DOI:** 10.1101/692673

**Authors:** Maddie E. James, Maria C. Melo, Federico Roda, Diana Bernal-Franco, Melanie J. Wilkinson, Greg M. Walter, Huanle Liu, Jan Engelstädter, Daniel Ortiz-Barrientos

**Affiliations:** The University of Queensland, School of the Environment, St Lucia, Queensland, Australia; Australian Research Council Centre of Excellence for Plant Success in Nature and Agriculture, The University of Queensland, St Lucia, Queensland, Australia; Universidad Nacional de Colombia, Departamento de Biología, Bogotá, Colombia; Los Andes University, Bogotá, Colombia; Queensland Alliance for Agriculture and Food Innovation, St Lucia, Queensland, Australia; University of Tasmania, School of Natural Sciences, Hobart, Australia; Centre for Genomic Regulation, Barcelona, Catalonia, Spain

**Keywords:** Parallel speciation, mutation-order, ecological speciation, natural selection, reproductive isolation, ecotypes

## Abstract

Natural selection shapes how new species arise, yet the mechanisms that generate reproductive barriers remain debated. Although ecological divergence in contrasting environments and mutation-order processes in similar environments are often viewed as distinct speciation mechanisms, we show they act together as part of a continuum we call ‘parallel mosaic speciation.’ In the *Senecio lautus* species complex, Dune and Headland ecotypes have evolved repeatedly along the Australian coastline. Through crossing experiments and field studies, we find that divergent natural selection promotes strong reproductive isolation between the Dune and Headland ecotypes. While uniform selection maintains reproductive compatibility among ecologically similar Dune populations, Headland populations have evolved reproductive barriers despite their convergent prostrate phenotypes, likely driven by adaptation to heterogeneous environments. To understand how habitat heterogeneity contributes to patterns of reproductive isolation, we extend previous theoretical work on the accumulation of hybrid incompatibilities to account for environmental gradients and polygenic adaptation. We show that the probability of reproductive isolation depends on three factors: how similar the environments are, how complex the genetic architecture is, and how selection coefficients are distributed among beneficial mutations. These theoretical findings explain how reproductive isolation arises in systems like *Senecio*, where multiple forms of selection jointly drive parallel speciation.

## INTRODUCTION

Speciation, the process by which new species arise, is a fundamental driver of biodiversity. However, demonstrating the direct role of natural selection in speciation remains a challenge. Systems exhibiting parallel evolution—where populations independently acquire similar phenotypes under comparable selective pressures—provide a powerful framework for investigating the role of selection in speciation (Schluter 2001; Langerhans and Riesch 2013). These natural replicates of the evolutionary process (Schluter and Nagel 1995; Schluter 2000) enable a thorough examination of the mechanisms that drive reproductive isolation. Current theoretical frameworks distinguish two primary modes of speciation by natural selection: ecological speciation, driven by deterministic adaptation to contrasting environments, and mutation-order speciation, where stochastic processes generate reproductive barriers between populations adapting to similar conditions (Schluter 2009; Schluter and Conte 2009; Nosil and Flaxman 2011; Nosil 2012; Crespi and Nosil 2013, but see Sobel et al. 2010; Langerhans and Riesch 2013).

Parallel ecological speciation occurs when divergent natural selection repeatedly generates reproductive isolation between populations adapting to contrasting environments, while maintaining reproductive compatibility among populations experiencing similar selective pressures (Figure 1A; Schluter and Nagel 1995; Ostevik et al. 2012). The resulting reproductive barriers can manifest through environment-dependent isolation, where immigrants and hybrids show reduced fitness in parental habitats, and environment-independent isolation, characterized by intrinsic hybrid incompatibilities regardless of ecological context (Rundle and Nosil 2005; Nosil 2012). The threespine stickleback (*Gasterosteus aculeatus*) system exemplifies this process, where the repeated colonization of freshwater habitats by marine ancestors has driven parallel evolution of adaptive traits, including reduced body armor and modified feeding morphology (Colosimo et al. 2005; Wund et al. 2008). Reproductive isolation between marine and freshwater forms has evolved primarily through extrinsic mechanisms, while geographically separated freshwater populations maintain reproductive compatibility (Rundle et al. 2000; McKinnon and Rundle 2002; McKinnon et al. 2004). This pattern—where ecological divergence predicts reproductive isolation more strongly than geographical separation—provides compelling evidence for the role of natural selection in speciation. Similar patterns of parallel ecological speciation have been documented in other animal taxa, including *Littorina* snails (Johannesson et al. 2010, 2024) and *Timema* walking stick insects (Nosil et al. 2002; Soria-Carrasco et al. 2014), although comparable evidence in plants remains limited (see Ostevik et al. 2012; James et al. 2023).

**Figure 1.**
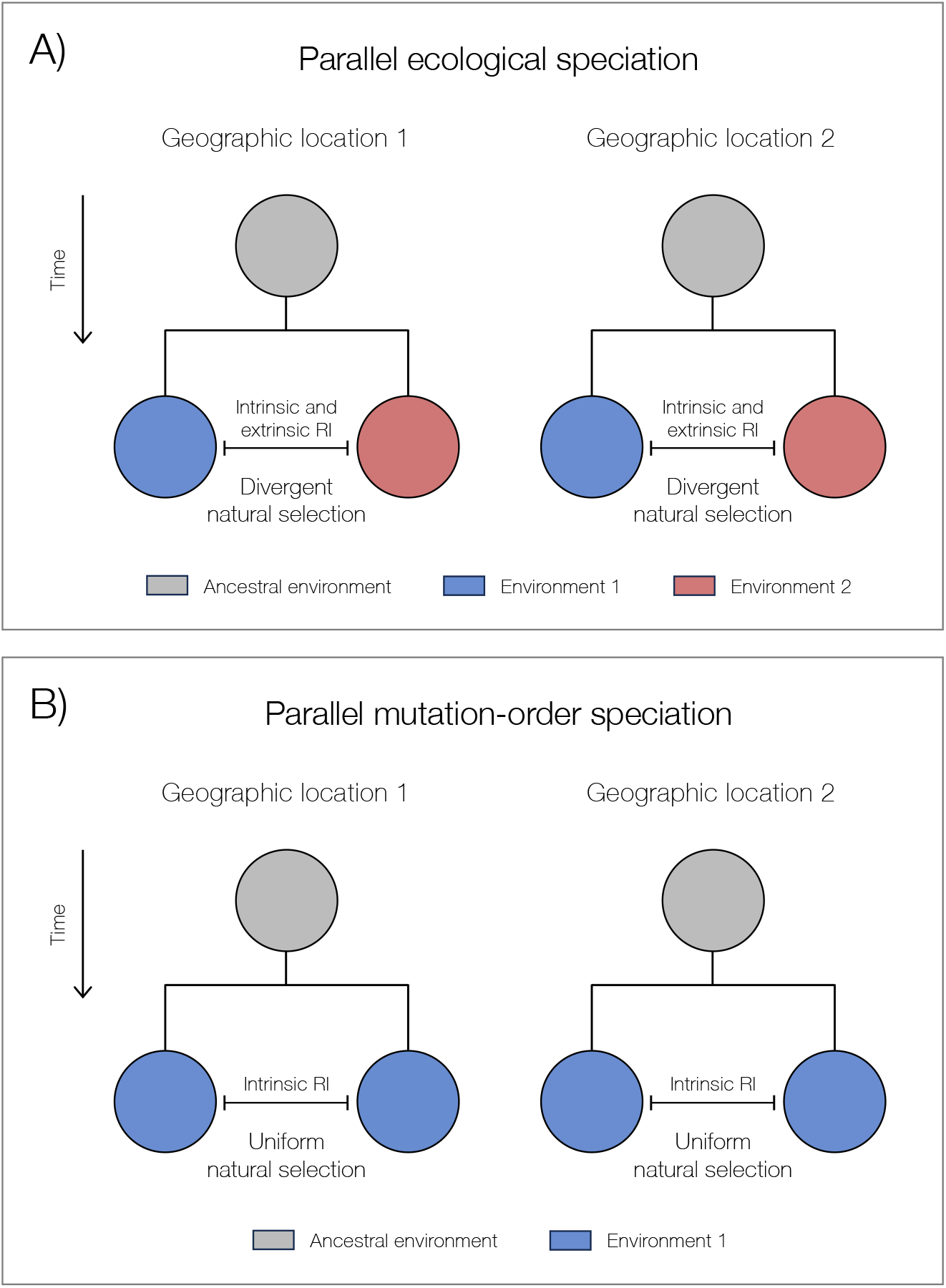
Schematic diagram representing the differences between parallel ecological speciation and mutation-order speciation. Circles represent populations, whereas colors represent different environmental conditions. Ancestral populations are in grey. A) In ecological speciation, intrinsic and extrinsic reproductive isolation (RI) evolves during the adaptation of populations to different environments (i.e., during divergent natural selection selection). B) In contrast, during mutation-order speciation, intrinsic reproductive isolation evolves during the adaptation of populations to similar environments (i.e., during uniform natural selection).

Mutation-order speciation is an alternative mechanism distinct from ecological divergence that can generate reproductive isolation. Under this process, uniform natural selection favors the same beneficial alleles across populations adapting to the same selective pressures (Figure 1B). However, reproductive incompatibilities arise as different beneficial alleles randomly arise and fix in each population. These alleles are incompatible when combined in hybrid genomes due to negative epistatic interactions, and as a result, can cause reproductive isolation between taxa (Mani and Clarke 1990; Schluter 2009; Nosil and Flaxman 2011; Nosil 2012). The mechanistic basis of mutation-order speciation differs fundamentally from ecological speciation: rather than arising from maladaptation to parental environments, reproductive barriers emerge from deleterious interactions between independently acquired beneficial mutations. Because these mutations experience positive selection, their fixation and subsequent incompatibilities can accumulate more rapidly than under neutral divergence through genetic drift (Mani and Clarke 1990).

Experimental studies provide compelling evidence for mutation-order speciation, demonstrating the evolution of reproductive barriers between populations adapting to identical environments such as in *Drosophila* (Hsu et al. 2024) and *Saccharomyces* (Ono et al. 2017). While the strictest definitions of mutation-order speciation require populations to inhabit identical environments (Mani and Clarke 1990), some definitions acknowledge it can also occur when populations adapt to similar but non-identical habitats (e.g., Schluter 2009; Schluter and Conte 2009; Nosil and Flaxman 2011). This makes documenting mutation-order speciation in nature challenging: determining whether reproductive barriers arose from the stochastic fixation of beneficial mutations or subtle ecological differences can blur the distinction between mutation-order and ecological processes. Addressing this challenge requires systematic investigation of speciation mechanisms across diverse taxa with broad geographic ranges (Anderson and Weir 2022), particularly in understudied groups such as plants.

The *Senecio lautus* species complex (Ali 1964; Radford et al. 2004; Thompson 2005) provides an exceptional system for examining the relationship between natural selection and speciation. Two phenotypically distinct ecotypes have repeatedly evolved in the species complex: an ancestral erect Dune ecotype occupying sheltered sand dunes and a derived prostrate Headland ecotype inhabiting exposed rocky outcrops (Figure 2A). These ecotypes form multiple Dune-Headland parapatric pairs along the Australian coastline (Figure 2B), which can be considered natural replicates of the evolutionary process (Roda et al. 2013a; James et al. 2021a). Each ecotype is locally adapted to a distinct habitat type (Roda et al. 2013b; Walter et al. 2016, 2018a; James et al. 2021b), yet Headland sites exhibit fine-scale heterogeneity in soil chemistry, microclimate, and biotic interactions, which generates a complex mosaic of selective pressures (Roda et al. 2013b; Walter et al. 2016). This environmental variation has significant implications for the genetic architecture of adaptation: while Headland populations exhibit convergent prostrate phenotypes, they achieve this convergence through distinct combinations of small-effect alleles, revealing the polygenic nature of adaptive evolution in the system (Roda et al. 2013b; James et al. 2021b, 2023a; Kaur 2024).

**Figure 2.**
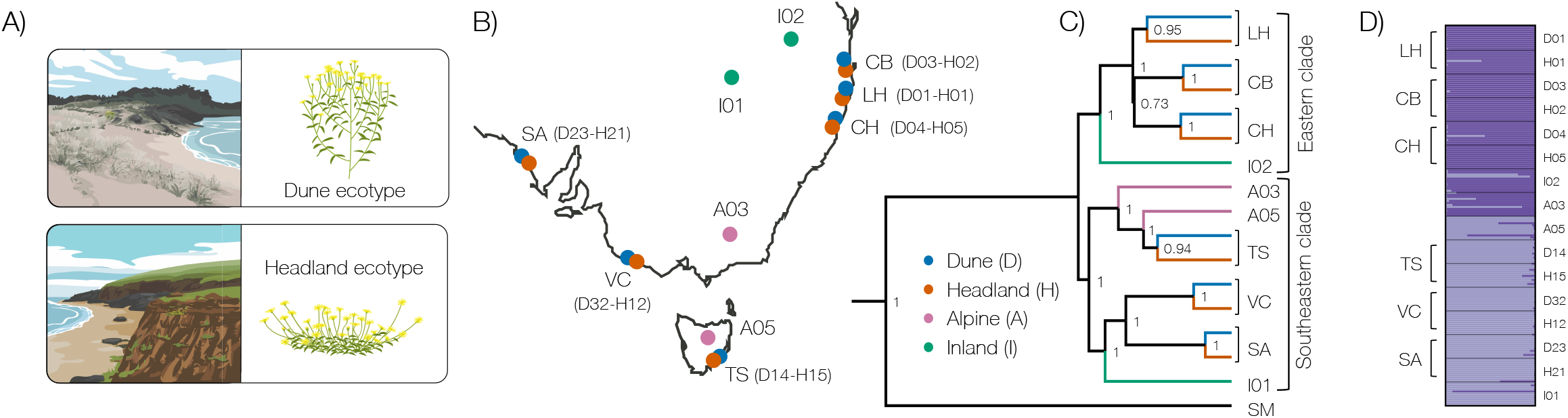
Parallel evolution of *Senecio lautus* Dune and Headland populations. A) Illustrations of the Dune and Headland environments and ecotypes. B) Geographic distribution in Australia of the 16 populations in the study (blue Dune ecotype, orange Headland ecotype, pink Alpine ecotype, green Inland ecotype). C) Bayesian phylogeny constructed with 13 neutral markers implemented in *BEAST. Numbers on nodes are credible posterior probabilities. D) Bayesian assignment of individuals to genetic clusters within STRUCTURE for *K* = 2. Each individual is depicted as a bar, with colors representing ancestry proportions to each K cluster.

Research in *Senecio* has revealed complex interactions between environment-dependent and intrinsic reproductive barriers. Reciprocal transplant experiments demonstrate extrinsic isolation between Dune and Headland ecotypes, indicating that divergent selection maintains reproductive boundaries through adaptation to contrasting environments (Melo et al. 2014; Richards and Ortiz-Barrientos 2016; Richards et al. 2016; Walter et al. 2016, 2018b; Wilkinson et al. 2021). While F_1_ hybrids between eastern coastal populations are largely reproductively compatible (Melo et al. 2014; Richards and Ortiz-Barrientos 2016; Walter et al. 2016), F_2_ and subsequent generations display strong forms of reproductive isolation (Walter et al. 2020; Wilkinson et al. 2021). Finally, the identification of a shared genetic basis between gravitropism—a key ecological trait differentiating the ecotypes—and hybrid sterility in *Senecio* demonstrates a direct pathway through which natural selection on ecological traits promotes speciation (Wilkinson et al. 2021).

In this work, we use empirical analyses to test fundamental predictions of parallel ecological and mutation-order speciation in the *Senecio lautus* species complex. We investigate how environmental heterogeneity shapes the accumulation of reproductive barriers and create a mathematical framework to explicitly incorporate environmental gradients and polygenic architectures, providing novel insights into speciation across heterogeneous landscapes. Our work shows that deterministic and stochastic processes interact in complex ways to generate reproductive isolation during parallel speciation.

## METHODS

### Phylogenetic Independence of Replicate Populations

#### Samples

We studied parallel speciation in six parapatric Dune-Headland *Senecio lautus* population pairs from the eastern and southern coasts of Australia (Figure 2A, Table S1). For each population, we used previously collected leaf samples from 12 individuals (Roda et al. 2013a). To strengthen our phylogenetic analysis, we included two additional ecotypes (two Inland and two Alpine populations; *n*_*pop*_ = 11), and the closely related African species *S. madagascariensis* (see Roda et al. 2013) as an outgroup (*n* = 4) (Figure 2A, Table S1). We extracted DNA from each sample using a modified CTAB protocol (Clarke 2009), purified samples using Promega’s Wizard® SV Gel and PCR Clean-Up System, and standardized each sample to 30ng/μL. We undertook targeted re-sequencing of nuclear genomic regions, resulting in 13 neutral markers across all populations (see *Supplementary Material 1* for details on primers, library preparation, sequencing, bioinformatics and neutrality tests; Table S2, S3). For a detailed taxonomy of the *S. lautus* species complex, please see supplementary tables in Roda et al. (2013a).

#### Phylogeny

Previous research in *S. lautus* demonstrated phylogenetic independence of five out of the six Dune-Headland population pairs in the current study (James et al. 2021a). To confirm the independence of all six pairs, we conducted a Bayesian phylogenetic analysis using the 13 neutral loci with *BEAST v1.7.5 (Heled and Drummond 2010), an extension of BEAST (Drummond et al. 2012) designed for multilocus and multi-individual species tree estimation. We generated the XML file for *BEAST using BEAUTi v1.7.5 (Drummond et al. 2012), and ran the analysis with a chain length of 300,000,000 under a strict molecular clock. Using ITS as our reference locus (mutation rate = 4.13×10^-9^ subs/site/year for herbaceous plants; Kay et al. 2006), we estimated relative mutation rates for all other loci. The HKY model (Hasegawa et al. 1985) best fit our 13 loci based on the Bayesian Information Criterion implemented in jModelTest (Posada and Crandall 1998). We used a Yules speciation process for species tree estimation, which assumes that lineages split at a constant rate. We generated the maximum clade credibility tree in TreeAnnotator v1.7.5 (Drummond et al. 2012) with a burn-in of 10,000 steps, and visualized the tree in FigTree v1.4.4 using *S. madagascariensis* as the outgroup.

#### Population Structure

We assessed population structure by identifying the number of genetic clusters (K) across all populations using STRUCTURE v2.3.4, a Bayesian Markov Chain Monte Carlo (MCMC) approach (Pritchard et al. 2000). We performed analyses with the variable sites of the 13 neutral loci using an admixture model and the correlated allele frequency model (Falush et al. 2003). Following guidelines from Gilbert et al. (2012) and Janes et al. (2017), we tested K values from 1 to 16, running 20 iterations per K with a burn-in of 100,000 and MCMC run length of 100,000. We confirmed convergence of model parameters through visual inspection of MCMC summary statistics. The most likely K value was evaluated using methods from Pritchard et al. (2000) and Evanno et al. (2005), implemented in STRUCTURE HARVESTER v0.6.93 (Earl and vonHoldt 2012). Since both methods tend to overestimate K, and higher K values provided no additional clustering information, we selected the smallest K that captured the major population structure in the data. We aligned cluster assignments across iterations using CLUMPP v1.1 (Jakobsson and Rosenberg 2007) with the complete search algorithm, and visualized results using DISTRUCT v1.1 (Rosenberg 2004).

### Intrinsic Reproductive Isolation Between and Within Ecotypes

#### Samples

To investigate patterns of reproductive isolation, we selected four Dune-Headland population pairs: two from the eastern clade (LH and CH) and two from the southeastern clade (VC and SA) of the phylogeny (Figure 2; Table S1). From each population, we collected seeds from 30 individuals in the field spaced at least ten meters apart. We stored seeds in dry conditions at 4°C at The University of Queensland. To create seed stocks for each population as well as to eliminate maternal effects (Bischoff and Müller-Schärer 2010), we first germinated and grew plants for one generation. To induce germination, we scarified seeds by trimming 1 mm off the micropyle side and placed them on moist filter paper in Petri dishes in a controlled growth room at 25°C. We kept seeds in darkness for three days to promote root elongation, and then transferred to a 12h:12h light:dark cycle for seven days to encourage vegetative growth. We then transplanted seedlings into 0.25L pots containing a soil mix (70% pine bark, 30% coco peat) supplemented with slow-release Osmocote fertilizer (5 kg/m^3^) and Suscon Maxi insecticide (830 g/m^3^). After two months, we conducted controlled crosses by repeatedly rubbing mature flower heads together over three to five days. This ensured bi-directional pollen transfer and maximum fertilization opportunity. The resulting seeds were stored at 4°C.

#### Reproductive Isolation Measures

We quantified two components of intrinsic reproductive isolation: F_1_ seed set and F_1_ viability (germination). To assess these barriers, we compared the success of crosses between populations of the *same* ecotype (D × D and H × H population crosses) with those of *different* ecotypes (D × H). This was undertaken both *within* and *between* the eastern and southern phylogenetic clades to account for phylogenetic divergence time (Coyne and Orr 1997). We also conducted intra-population crosses as controls. Using the protocol outlined above, we grew up to 14 families from each of the eight populations from the eastern and southeastern clades of the phylogeny (Table S1). When the plants flowered, we performed intra- and inter-population crosses twice daily, completing 260 crosses in total (see Table S4).

To measure reproductive isolation at the F_1_ seed set stage, we counted the proportion of fertilized seeds per flower head, identifying unfertilized seeds by their thin size and pale color. We quantified the strength of reproductive isolation (RI) for F_1_ seed set between populations as:

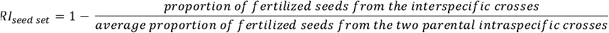

To assess reproductive isolation at the F_1_ viability stage, we measured germination rates from the crosses defined above. For each of the 260 crosses, we germinated five seeds by placing them onto moist filter paper in Petri dishes (one cross per Petri dish). We used the same germination conditions as above but without scarification to test the intrinsic ability of embryos to emerge from the seed coat. The strength of reproductive isolation for F_1_ viability was calculated as:

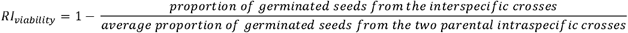

#### Statistical Analyses

We analyzed patterns of intrinsic reproductive isolation using linear mixed effect models in R v4.1.0 (R Core Team 2021) with the *lmerTest* package (Kuznetsova et al. 2017):

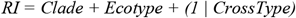

where *RI* is the reproductive isolation values for either F_1_ seed set or F_1_ viability, *Clade* indicates whether crosses occurred within or between phylogenetic clades, *Ecotype* denotes whether crosses were between the same or different ecotypes, and *CrossType* is a random effect with a random intercept model, representing the population comparison. For both F_1_ seed set and viability models, we initially tested for an interaction between *Clade* and *Ecotype* but removed it due to non-significance. To examine specific cross-type effects, we ran additional linear models:

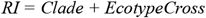

where *EcotypeCross* specifies the exact cross comparison (D × D, H × H, D × H, or H × D). Again, we removed the interaction terms as they were non-significant. Finally, we used one-sided t-tests to determine whether reproductive isolation values for D × D and H × H crosses (both within and between clades) were significantly greater than zero.

### Extrinsic Reproductive Isolation Within Ecotypes

#### Samples

We tested for mutation-order effects on speciation by examining extrinsic reproductive isolation within ecotypes. If this type of speciation is occurring in *Senecio*, populations of the same ecotype should exhibit similar performance at other localities within the same environment. This was assessed by reanalyzing data from a previous field transplant experiment by Walter et al. (2016). Briefly, in this study seeds of six populations (three Dune-Headland replicate pairs) were generated under common garden conditions and then transplanted (n = 150-180 seeds per population in each environment; Table S1) into the dune and headland environments of the Lennox Head (LH) population pair. Seeds were planted in fully randomized grids with six replicate blocks in each environment, seedling survival was monitored over time, and seedling establishment was recorded when plants had grown ten true leaves. For further details, see Walter et al. (2016).

#### Statistical Analyses

We compared local population performance (LH Dune seeds planted in the LH dune habitat; LH Headland seeds planted in LH headland habitat) against non-local populations (i.e., Stradbroke Island and Cabarita Beach Dune and Headland seeds in the LH dune and headland environments, respectively). Our analyses examined two fitness components: survival and establishment.

Using the R package *coxme* (Therneau 2024), we modeled survival separately for dune and headland environments:

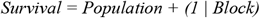

where *Survival* is days survived (censored at day 320), *Population* indicates seed source location, and *Block* is a random effect for replicate experimental blocks within each environment.

We then tested whether local populations showed similar probabilities of reaching seedling establishment compared to non-local populations using a generalized linear mixed-effects model in the R package *lme4* (Bates et al. 2015). The model used the binomial family with a logit link function to model the probability of establishment success. We analyzed dune and headland environments separately, with the model:

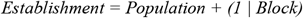

where *Establishment* is a binary variable indicating whether a plant reached ten leaves and *Block* is a random effect for replicate experimental blocks within each environment.

In both analyses, we tested whether non-local populations performed similarly to the local population. To do this, we explicitly set the local population as the reference level (intercept) for the *Population* factor. This parameterization allows coefficient estimates to represent deviations of non-local populations from local population performance, where significant negative coefficients indicate local adaptation.

### Connecting Ecology and Phenotype with Intrinsic Reproductive Isolation

To further assess the role of mutation order in driving parallel speciation in *Senecio*, we examined the relationship between intrinsic reproductive isolation and environmental or phenotypic distance. For mutation-order to hold true, we expect no associations because populations are expected to fix advantageous mutations randomly rather than in response to ecological or phenotypic differences (Schluter 2009).

#### Intrinsic Reproductive Isolation and Environmental Distance

To test the role of ecological differences to intrinsic isolation, we explored the association between intrinsic reproductive isolation (F_1_ seed set and F_1_ viability, as measured above) and environmental variance among Headland populations. We used previously published soil data from Roda et al. (2013b) to quantify environmental variance, where 38 soil variables (nutrients, salts and metals) were measured for each population. We used R for statistical analyses and scaled each soil variable to a mean of zero and standard deviation of one. We used the *vegan* package (Blanchet et al. 2018) in R to create a distance matrix of the soil variables for each population. We performed multidimensional scaling to calculate the Euclidian distance between populations in environmental space using the first five principal component axes, which explained more than 95% of the variance. To estimate genetic distance, we used the *ape* package (Paradis et al. 2004) in R to calculate pairwise distances between populations based on branch lengths from the phylogeny constructed above.

To assess the relationship between reproductive isolation and environmental distances, we performed linear models:

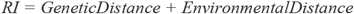

where *RI* is the reproductive isolation value for either F_1_ seed set or F_1_ viability.

#### Intrinsic Reproductive Isolation and Phenotypic Distance

We examined the association between intrinsic reproductive isolation (F_1_ seed set and F_1_ viability) with phenotypic variance between Headland populations. We used a combination of previously published phenotypic data from Walter et al. (2018a), and unpublished data for additional populations measured at the same time, where phenotypes were measured in controlled glasshouse conditions. This dataset consists of four plant architecture traits (vegetative height, stem length/width, number of branches, and stem diameter) and six leaf traits (area, perimeter^2^/area^2^, circularity, number of indents, indent width, and indent depth), measured for up to 17 individuals per population (Table S1), see Walter et al. (2018a) for specific details on growing conditions and trait measurements. As in the environmental analysis, we scaled each phenotypic variable to have a mean of zero and standard deviation of one, calculated the distance matrix of the phenotypic traits for each population, and performed multidimensional scaling to calculate the Euclidian distance between populations in environmental space using the first seven principal component axes, which explained more than 95% of the variance. We used the genetic distances calculated above in *ape*. We performed linear models to assess the relationship between reproductive isolation and phenotypic distances, again averaging the reciprocal crosses:

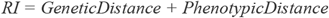

where *RI* is the reproductive isolation value for either the F_1_ seed set or F_1_ viability for each population comparison.

### Mathematical Analysis of Parallel Speciation

To further understand how various forms of natural selection drive patterns of reproductive isolation in *Senecio*, we created a mathematical framework to explore how both ecological and mutation-order processes can jointly underlie patterns of reproductive isolation. Our framework begins with the classic Dobzhansky-Muller model, which establishes how hybrid fitness decreases through epistatic interactions between incompatible alleles (Orr 1995). Next, we build a model based on the Unckless-Orr framework (Unckless and Orr 2009), which quantifies the probability of Dobzhansky-Muller Incompatibility (DMI) formation between populations adapting to the same environments – representing the strictest form of mutation-order speciation, where environmental conditions are identical between populations. To capture the dynamics of divergent natural selection, we extend the model to incorporate environmental heterogeneity, where DMIs accumulate across a gradient of environmental similarity. This gradient ranges from identical environments where mutation-order processes dominate, to contrasting environments where ecological divergence occurs. Additionally, we integrate polygenic adaptation into the framework, providing a more nuanced understanding of how incompatibilities evolve with increasing genetic complexity.

#### Foundations

The foundation of our analysis builds upon the classic Unckless-Orr model examining DMI formation between two populations adapting to identical environments (Unckless and Orr 2009). This model considers two interacting loci A and B, each with ancestral (*A*_0_, *B*_0_) and derived (*A*_1_, *B*_1_) alleles. The ancestral genotype (*A*_0_, *B*_0_) has a fitness of 1, while genotypes with single derived alleles *A*_1_, *B*_0_ and *A*_0_, *B*_1_ have fitness values of 1+ *s*_*A*_ and 1+ *s*_*B*_ respectively. The combination of both derived alleles (*A*_1_, *B*_1_) creates an incompatibility with fitness 1−*t*, where *t* quantifies the severity of the DMI (Orr 1995). When *t*= 1, the hybrid combination is completely inviable/sterile, while smaller values of *t* represent partial reproductive isolation. The inclusion of *t* in the model helps capture the concept that while individual-derived alleles can be beneficial in their genetic backgrounds, their combination in hybrids can be deleterious (*t* > 0). This is a key feature of the Dobzhansky-Muller model of speciation—incompatibilities arise not from individual mutations being deleterious, but from their negative epistatic interactions when brought together in hybrids.

Under strong selection (*Ns* >> 1) and weak mutation (*N*_*μ*_ ≪ 1), where adaptive trajectories retain stochastic elements despite directional selection, the probability of DMI formation depends on selection coefficients through the Unckless-Orr model:

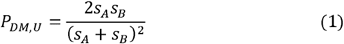

We use the subscript *U* to refer to the Unckless-Orr model. This equation reveals that during adaptation to identical environments, DMIs are most probable when selection coefficients are similar (*s*_*A*_ ≈*s*_*B*_), as populations experience maximum stochasticity in the order of mutation fixation. When selection coefficients are equal, either beneficial mutation is equally likely to fix first in each population, maximizing the probability that populations will fix different alleles. However, when selection coefficients differ substantially, both populations are likely to fix the mutation with the stronger selective advantage first, reducing the probability of DMI formation.

### Extensions

We extend this framework by modeling how environmental differences create asymmetric selection on alleles between populations. For any locus, we assume its selection coefficient in its “home” environment (*s*_*i*_) is reduced by a factor *φ* when it occurs in the alternative environment (*φs*_*i*_). This parameter *φ∈ [0,1]* captures the degree to which selection pressures transfer between environments, with *φ= 1* indicating identical selection and *φ= 0* indicating complete asymmetry where alleles that are beneficial in one environment are neutral in alternative environments.

Under these assumptions, the probability of DMI formation becomes:

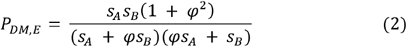

When environments are identical (*φ*= 1), the equation reduces to the classic Unckless-Orr model. As environmental differences increase (decreasing *φ*), selection becomes increasingly asymmetric between populations, making it more likely they will fix different alleles. In the extreme case of complete asymmetry (*φ*= 0), DMI formation becomes inevitable because alleles strongly favored in one environment are neutral in the other.

For polygenic traits involving *n* interacting loci with selection coefficients *s*_*i*_, the probability of at least one DMI forming is:

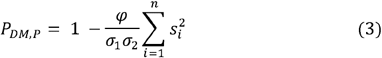

where σ_l_ and σ_2_ represent the sums of selection coefficients in each population. When environments are identical (*φ*= 1), this reduces to an *n*-locus extension of the two-locus Unckless and Orr (2010) model, while complete asymmetry (*φ*= 0) guarantees DMI formation. For equal selection coefficients (*s*_*i*_ = *s*), the formula simplifies to:

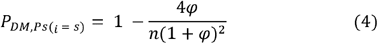

This shows how genetic complexity amplifies the effects of selection asymmetry through combinatorial growth in potential incompatibilities (see *Supplementary Material 2* for detailed derivations of all equations).

#### Implementation

We implemented these models using Python v3.10.9 and NumPy v1.23.5 (Harris et al. 2020) for numerical calculations, and Matplotlib v3.9.2 (Hunter 2007) for visualization. Our analysis explored selection coefficient space (*s*_*A*_ *υs s*_*B*_), environmental similarity (*φ*), and the impact of interacting loci number (*n*) on DMI probability, providing a theoretical explanation for the observed patterns of reproductive isolation in *Senecio*.

## RESULTS AND DISCUSSION

### Empirical Patterns of Parallel Speciation

#### Phylogenetic Independence Reveals Multiple Origins of Coastal Ecotypes

To investigate parallel speciation in *Senecio*, we first confirmed the independent origins of Dune and Headland populations in this study (also see Roda et al. 2013a; James et al. 2021a). Using 13 neutral markers, we found that populations cluster into two main clades (eastern and southeastern; Figure 2C, 2D), that align with their coastal geographic distribution (Figure 2B). As expected, the phylogenetic relationships support parallel evolution: Dune-Headland population pairs form sister taxa, with both ecotypes present within each clade (Figure 2C). This result confirms previous independent datasets in the *Senecio* system (Roda et al. 2013a; James et al. 2021a). Although gene flow can create false signals of parallel evolution (Endler 1977; Barton and Hewitt 1985; Coyne and Orr 2004; Bierne et al. 2013), previous work in *Senecio* showed that gene flow among *Senecio* populations, both within and between ecotypes, is negligible (James et al. 2021a). These findings indicate that the observed phylogenetic patterns reflect the independent adaptive divergence of Dune-Headland pairs along the Australian coastline.

### Divergent Selection Drives Strong Reproductive Isolation Between Ecotypes

Parallel ecological speciation generates specific predictions about patterns of reproductive isolation: populations adapting to contrasting environments should develop reproductive incompatibilities, while those inhabiting similar environments should maintain reproductive compatibility. In contrast, mutation-order processes predict that populations evolve reproductive barriers even when adapting to similar environments through the independent fixation of different beneficial alleles (Schluter and Nagel 1995; Ostevik et al. 2012). To evaluate these contrasting theoretical predictions in *Senecio*, we quantified intrinsic reproductive isolation between populations from the same environments (within Dune or within Headland ecotypes) and different environments (between Dune and Headland ecotypes). Our analysis focused on two quantitative components of reproductive isolation, measured relative to parental averages: F_1_ hybrid seed production and F_1_ hybrid viability during germination.

Consistent with the role of divergent selection, populations from similar environments showed higher reproductive compatibility than those from different environments. This pattern held for both seed set (*F*_l,15.73_ = 5.93, *P* = 0.023, *R*^*2*^ = 0.58 after taking into account the effect of clade *F*_l,l5. 61_ = 13.49, *P* = 0.002) and viability (*F*_1,14.48_ = 7.25, *P* = 0.017, *R*^2^ = 0.36 after taking into account the impact of clade *F*_1,14.48_ = 7.21, *P* = 0.018) (Figure 3A). Crosses within clades were generally viable, with crosses between the same ecotypes showing some hybrid vigor (reproductive isolation values < 0), suggesting complementary genetic interactions. The most substantial (almost complete) reproductive isolation occurred in Dune-Headland crosses between the two clades (Figure 3A). These results indicate that natural selection drives speciation in *Senecio*, with reproductive isolation increasing as genetic divergence accumulates between ecotypes. This pattern mirrors findings in other systems, including stickleback fish (Hendry et al. 2009; Stuart et al. 2017), *Littorina* snails (Johannesson et al. 2024), and cichlid fishes (Weber et al. 2021).

**Figure 3.**
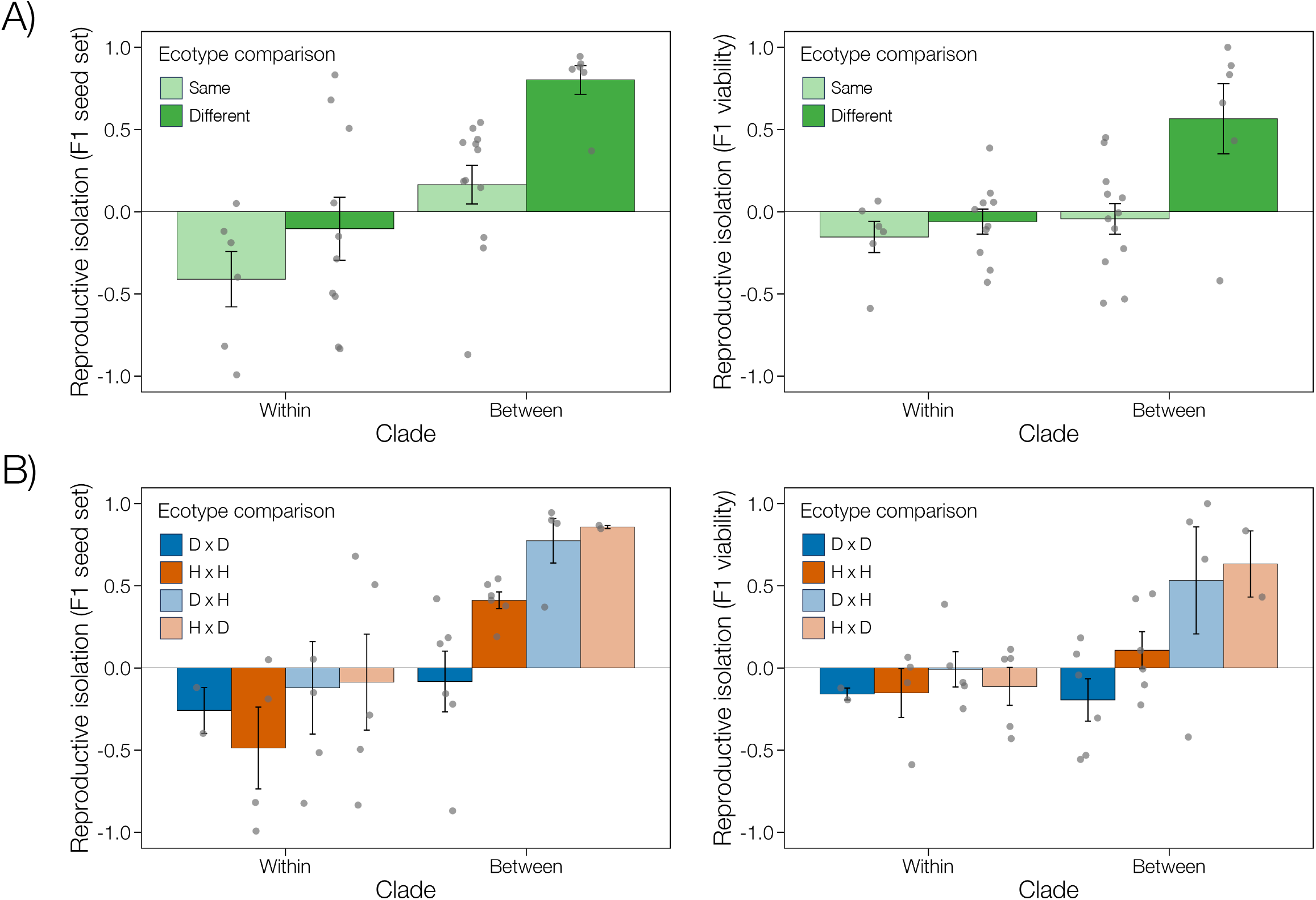
Strength of intrinsic reproductive isolation (RI) of coastal Dune (D) and Headland (H) *Senecio lautus* populations for F_1_ seed set (left graphs) and F_1_ viability. Data points represent the mean of multiple crosses for each population comparison (F_1_ seed set: n_total_ = 260 crosses; F_1_ viability: n_total_ = 259 crosses). Crosses were undertaken within and between the two clades defined in the phylogeny of Figure 1. Reproductive isolation was examined (A) between the same ecotype (D × D and H x H) or between different ecotypes (D × H and H × D), as well as (B) for each separate cross type (D × D, H × H, D × H and H × D). Positive values imply that hybrids perform worse than parents, and negative values that hybrids perform better than parents.

#### Unexpected Reproductive Barriers Emerge Among Convergent Headland Populations

To establish whether the observed patterns of intrinsic reproductive isolation represent parallel ecological speciation in *Senecio*, we must demonstrate reproductive compatibility within each ecotype, for both hybrid viability and fertility. Cross type (D × D, H × H, D × H, or H × D) significantly affected reproductive isolation for both F_1_ seed set (*F*_3,29_ = 3.44, *P* = 0.03, *R*^2^ = 0.46 after taking into account the effect of clade *F*_1,29_ = 14.44, *P* = 0.0007) and F_1_ viability (*F*_3,29_ = 3.52, *P* = 0.03, *R*^2^ = 0.34 after taking into account the effect of clade *F*_1,29_ = 4.51, *P* = 0.042) (Figure 3B). Dune populations showed the expected pattern of parallel ecological speciation, maintaining reproductive compatibility within and between clades. Their reproductive isolation values were not significantly greater than zero for either F_1_ seed set (within clades *t*_5_ = −1.85, *P* = 0.84, between clades *t*_5_ = −0.44, *P* = 0.66), or F_1_ viability (within clades *t*_5_ = 4.4, *P* = 0.93, between clades *t*_5_ = −1.51, *P* = 0.9). Headland populations were similarly compatible within clades for both F_1_ seed set (*t*_3_ = −1.96, *P* = 0.93) and F_1_ viability (*t*_3_ = 1.02, *P* = 0.81), and between clades for viability (*t*_5_ = −0.95, *P* = 0.19).

Our analyses revealed an unexpected pattern of reproductive incompatibility among Headland populations from different clades, manifested as significantly reduced F_1_ seed set (*t*_5_ = 8.12, *P* = 0.0002) (Figure 3B). While this pattern contradicts predictions of ecological speciation, it aligns with the expectation that mutation-order processes contribute to speciation. The reproductive isolation between Headland populations suggests parallel evolution of prostrate phenotypes occurred through distinct adaptive trajectories, resulting in different combinations of beneficial alleles in each population that create intrinsically unfit hybrids between populations. This interpretation is supported by previous genetic analyses in *Senecio*, which demonstrated that convergent Headland morphology evolved through the selection of distinct alleles and genes across populations (Roda et al. 2013b; James et al. 2021b). To further explore the role of mutation-order effects on speciation in *Senecio*, we tested additional predictions of speciation theory.

#### Ecological Differences do not Explain Reproductive Isolation in Headland Populations

Previous work in *Senecio* revealed contrasting patterns of environmental variation. Each ecotype occupies a distinct soil type, where variation within ecotypes is lower than between ecotypes (Roda et al. 2013b; Walter et al. 2016). The soils of different Headland populations share fundamental characteristics of being shallow and nutrient-rich but vary in specific mineral composition based on local geology. This habitat heterogeneity contrasts with the Dune populations, which occupy remarkably similar habitats even when separated by thousands of kilometers. To determine whether ecological variance reflect fitness variation within each ecotype, we reanalyzed reciprocal transplant experiments from Walter et al. (2016). In these experiments, three eastern-coast Dune populations were transplanted into the dune habitat at Lennox Head, while three eastern-coast Headland populations were transplanted into the rocky headland. Dune populations showed no signal of local adaptation: non-local individuals performed equally well or better than local Lennox Head individuals in both survival (Figure 4A) and establishment (Figure 4B). In contrast, the local Lennox Head Headland population consistently outperformed non-local populations in both fitness components (Figure 4).

**Figure 4.**
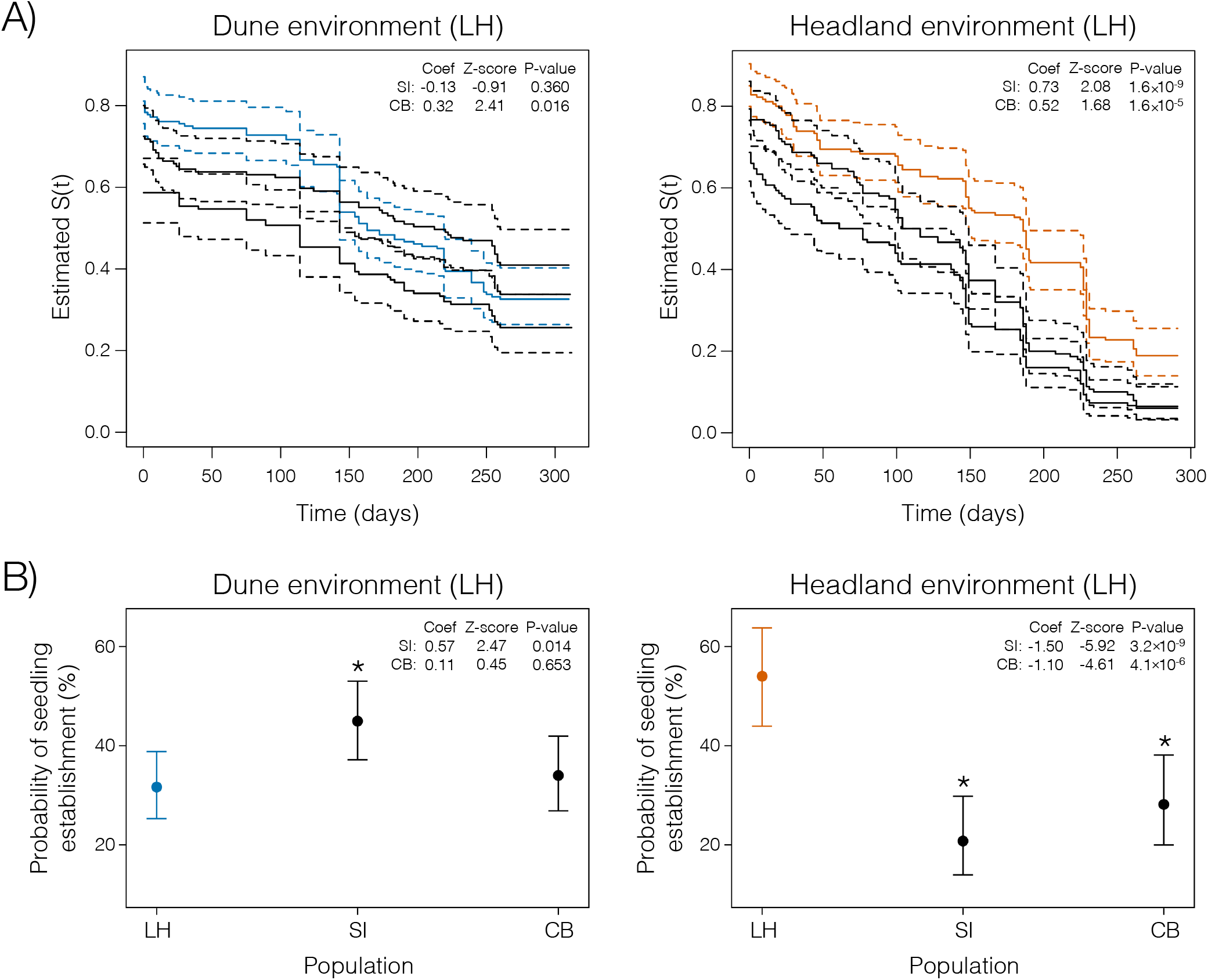
Strength of extrinsic reproductive isolation of coastal Dune and Headland *Senecio lautus* populations. Fitness of three replicate populations (LH – Lennox Head, SI – Stradbroke Island; CB – Cabarita Beach) from the Dune and Headland ecotypes transplanted into the dune and headland environments, respectively, at Lennox Head (Dune: n_total_ = 479 individuals; Headland: n_total_ = 480 crosses). Fitness was measured by A) survival curves, where the local population is denoted in color and dashed lines are 95% confidence intervals, and B) the probability of seedling establishment (production of 10 leaves), where asterisks denote significant differences from the local population.

Given the fitness differences among Headland populations at Lennox Head, we tested whether environmental variation (measured by soil ecology) explains patterns of intrinsic reproductive isolation across geography. If ecological divergence in soil composition drives reproductive barriers, we expect to find a correlation between measured environmental differences and intrinsic reproductive isolation across Headland populations. After controlling for genetic distance, we found no significant relationship between soil-based ecological divergence and reproductive isolation for F_1_ seed set (*F*_1,2_ = 3.00, *P* = 0.228), or F_1_ viability (*F*_1,2_ = 0.16, *P* = 0.729). The absence of an environment-reproductive isolation correlation for soil composition is consistent with mutation-order effects on speciation, where populations fix advantageous mutations randomly rather than in response to ecological differences (Schluter 2009). However, we cannot disregard the potential role of ecological differences in generating reproductive incompatibilities for two main reasons: other non-measured ecological variables may be the main selective force driving the association, and the relatively small sample size of soil data reduces our power to detect an effect.

#### Phenotypic Divergence does not Predict Reproductive Isolation in Headland Populations

We next asked whether subtle phenotypic variation among Headland populations could explain their reproductive isolation patterns. If divergent rather than uniform selection drives intrinsic reproductive isolation between Headlands, populations with greater phenotypic differences should show stronger reproductive barriers. However, after controlling for genetic distance, we found no significant relationship between phenotypic divergence and reproductive isolation for either F_1_ seed set (*F*_l,2_ = 12.88, *P* = 0.070) or F_1_ viability (*F*_l,2_ = 4.45, *P* = 0.170). The absence of a correlation between phenotypic divergence and intrinsic reproductive isolation further supports mutation-order effects on speciation, where genetic incompatibilities independently arise and contribute to the same adaptive phenotypes across populations (Schluter 2009). However, we again cannot disregard a possible association between phenotypic differences and reproductive incompatibilities due to potential unmeasured ecological variables and relatively small sample sizes for this analysis.

### Parallel Mosaic Speciation in *Senecio*

The patterns of reproductive isolation in the *Senecio lautus* species complex present an evolutionary paradox that challenges our traditional understanding of parallel speciation (Figure 6). While divergent natural selection in contrasting coastal environments has repeatedly driven the evolution of reproductive barriers between Dune and Headland ecotypes (Melo et al. 2014; Richards and Ortiz-Barrientos 2016; Richards et al. 2016; Walter et al. 2016, 2018b; Wilkinson et al. 2021), the pattern within each ecotype reveals an unexpected asymmetry. In Dunes, populations experience relatively uniform selection due to consistent environmental pressures (Roda et al. 2013b; Walter et al. 2016), allowing for reproductive compatibility despite geographic separation of thousands of kilometers and ancient divergence. Crosses between distant Dune populations even exhibit enhanced fitness through hybrid vigor. In contrast, Headland populations have evolved reproductive barriers in crosses between the phylogenetic clades, despite convergence on similar prostrate phenotypes. This suggests that the repeated evolution of similar phenotypes may obscure profound differences in how populations traverse adaptive landscapes. These incompatibilities between Headlands do not correlate with environmental or morphological differences, suggesting that mutation-order processes are at play. However, environmental heterogeneity (Roda et al. 2013b; Walter et al. 2016) and localized selective pressures among Headland populations indicate that patterns of reproductive isolation do not strictly align with mutation-order speciation.

In *Senecio*, this combination of divergent, uniform and heterogeneous selection forms what we call ‘parallel mosaic speciation’––a continuum of speciation patterns where reproductive isolation arises through multiple mechanisms (also see Langerhans and Riesch 2013). While theoretical frameworks have traditionally treated mutation-order and ecological speciation as distinct processes (Schluter 2009; Nosil and Flaxman 2011), empirical evidence suggests natural systems operate along a continuum of selective pressures (Johannesson et al. 2024). This complexity is particularly evident in plants, such as the *Senecio* system, where fine-scale environmental heterogeneity creates intricate selective mosaics that challenge the binary classification of uniform versus divergent selection (Lowry et al. 2019; James et al. 2023b). Here our parallel mosaic speciation framework explicitly incorporates within-ecotypic environmental heterogeneity to illustrate how various forms of natural selection shape patterns of reproductive isolation across heterogeneous landscapes.

Parallel mosaic speciation creates several theoretical challenges: How do systems of parallel evolution evolve different forms of reproductive isolation within and between ecotypes? How can populations that evolved nearly identical phenotypes accumulate reproductive barriers? And what is the effect of genetic architecture and environmental variance in creating patterns of reproductive isolation during parallel speciation? Addressing these questions requires an integrated theoretical framework to explain parallel speciation under mixed selection regimes and complex genetic architectures. Although theory has addressed components of these problems (e.g., Orr 1995; Barton 2001; Orr and Turelli 2001; Porter and Johnson 2002; Gavrilets 2004; Palmer and Feldman 2009; Fierst and Hansen 2010; Nosil and Flaxman 2011; Bank et al. 2012; Chevin et al. 2014; Fraïsse et al. 2014; Yamaguchi and Otto 2020; Thompson et al. 2024), many models oversimplify the genetic architectures underlying adaptation. Additionally, current frameworks inadequately account for how genetic architecture interacts with environmental heterogeneity to drive the accumulation of reproductive barriers. By integrating these factors, we can better understand how speciation unfolds across heterogeneous landscapes, moving us beyond the simple dichotomy of ecological vs. mutation-order speciation.

### Environmental Symmetry Determines the Probability of Incompatibility Formation

We developed a mathematical framework demonstrating how environmental symmetry and genetic architecture shape reproductive isolation during parallel mosaic speciation (Table 1). We found that the probability of Dobzhansky-Muller Incompatibility (DMI) formation is governed by the interaction of three key parameters: the distribution of selection coefficients (s), environmental symmetry (*φ*), and the genetic architecture (*n*) of the interacting loci. The classic Unckless-Orr model (*P*_*DM*, *U*_) demonstrates that under identical environments, the probability of DMI formation reaches a maximum when selection coefficients are equal (Table 2). This upper bound reflects maximum stochasticity in adaptive trajectories: when mutations confer equal fitness benefits (*s*_*A*_ = *s*_*B*_), either allele is equally likely to fix first in separate populations, maximizing the probability of incompatible combinations.

**Table 1.** Theoretical framework for parallel mosaic speciation. Dobzhansky-Muller Incompatibilities (DMIs) accumulate between populations through asymmetric selection effects and genetic architecture. The framework extends classic models to incorporate environmental heterogeneity and polygenic adaptation, providing a quantitative foundation for understanding parallel mosaic speciation.

| Component | Mathematical Form | Key Parameters |
| --- | --- | --- |
| Classic Unckless-Orr Model | $P_{DM,U} = (2s_A s_B) / (s_A + s_B)^2$ | $s_A, s_B$: selection coefficients |
| Environmental Effects Model | $P_{DM,E} = \frac{s_A s_B (1 + \varphi^2)}{(s_A + \varphi s_B)(\varphi s_A + s_B)}$ | $\varphi$: environmental symmetry (0 to 1) |
| Polygenic Architecture Model | $P_{DM,P} = 1 - \frac{\varphi}{\sigma_1 \sigma_2} \sum_{i=1}^n s_i^2$ | $n$: number of interacting loci |

**Table 2.** Unification of speciation models through a parallel mosaic framework. Environmental symmetry (*φ*) and selection coefficients (*s*) interact to produce diverse patterns of reproductive isolation across heterogeneous landscapes. Classic parallel ecological and mutation-order speciation represent extreme cases of parallel mosaic speciation where environmental heterogeneity and genetic architecture jointly determine the probability of speciation. *φ* refers to how selection coefficients transfer between environments. For instance, high symmetry means an allele experiences similar selection pressure in both populations. Also see Table 1.

| Model Component | Environmental Symmetry ($\varphi$) | Selection Coefficients | Population Structure | Reproductive Isolation | Example Systems |
| --- | --- | --- | --- | --- | --- |
| Parallel Ecological Speciation | $\varphi = 0$ between environments<br>$\varphi = 1$ within environments | $s_A \gg s_B$<br>(highly asymmetric) | Discrete habitat transitions | Strong between environments;<br>absent within environments | Threespine stickleback<br>marine-freshwater pairs<br>(Rundle et al. 2000) |
| Mutation-Order Speciation | $\varphi = 1$<br>(symmetric selection) | $s_A \approx s_B$<br>(similar magnitudes) | Independent populations<br>in similar habitats | Present despite selection<br>symmetry | Experimental<br>*Drosophila* lineages<br>(Hsu et al. 2024) |
| Parallel Mosaic Speciation | Continuous: $0 \leq \varphi \leq 1$ | Variable relationship<br>between $s_A$ and $s_B$ | Spatially structured<br>populations across<br>environmental mosaics | Varies with $\Phi$ and selection<br>coefficient relationships | *Senecio lautus* complex |

### The Role of the Environment in Mosaic Speciation

We extended the Unckless-Orr model by incorporating environmental symmetry (*φ*), to reveal how environmental heterogeneity systematically modifies DMI formation probability through its effects on relative selection pressures (Figure 5A). This environmental similarity parameter enables us to model speciation scenarios beyond the binary categorisation of identical versus contrasting environments, offering a more nuanced perspective on how environmental gradients drive reproductive isolation. This extended model (Equation 2, *P*_*DM*, *E*_) implies the following. First, when the environments are identical between populations (*φ* = 1), *P*_*DM*, *E*_ reduces to *P*_*DM*, *U*_, recovering classic predictions about the relationship between selection coefficients and reproductive isolation. This is akin to the *Senecio* Dune populations, where populations remain reproductively compatible due to nearly identical selective pressures. Second, as environments become increasingly different between populations (*φ* → 0), the probability of DMI formation approaches unity regardless of selection coefficient ratios, demonstrating how environmental differences can override deterministic selection effects, as seen between the Dune and Headland ecotypes. Third, for intermediate symmetry (0 < *φ* < 1), the probability of DMI formation increases monotonically with decreasing *φ* while becoming progressively less dependent on the ratio *s*_*B*_*/s*_*A*_. This is similar to the Headland populations, which experience some degree of environmental heterogeneity.

**Figure 5.**
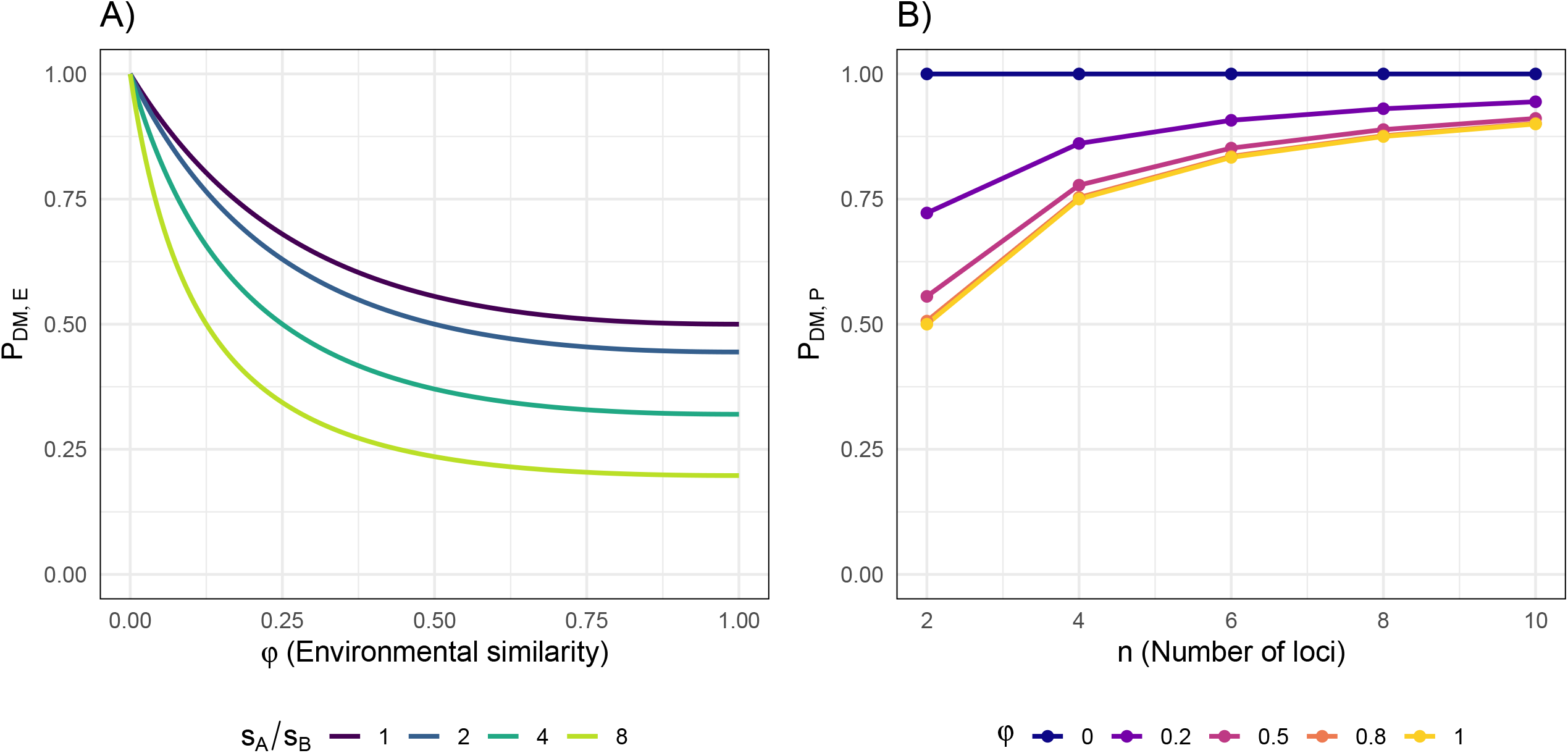
Environmental and polygenic effects on Dobzhansky-Muller incompatibility (DMI) formation probability. A) Environmental effects model (*P*_*DM*, *E*_), showing how the probability of incompatibility formation varies with environmental similarity *φ* under different selection ratios *s*_*A*_*/s*_*B*_. Lower *φ* (greater environmental difference) increases DMI probability, particularly when selection is more asymmetric (larger *s*_*A*_*/s*_*B*_). In the boundary case *φ*= 1 and *s*_*A*_ = *s*_*B*_, this single□locus model reduces to the Unckless–Orr result of *P*_*DM*, *U*_ = 0.5 (purple line). B) Polygenic architecture model (*P*_*DM*, *P*_), illustrating how the probability of DMI increases with the number of contributing loci *n*. Genetic complexity amplifies the impact of selection on reproductive isolation, especially under strongly asymmetric selection regimes.

**Figure 6.**
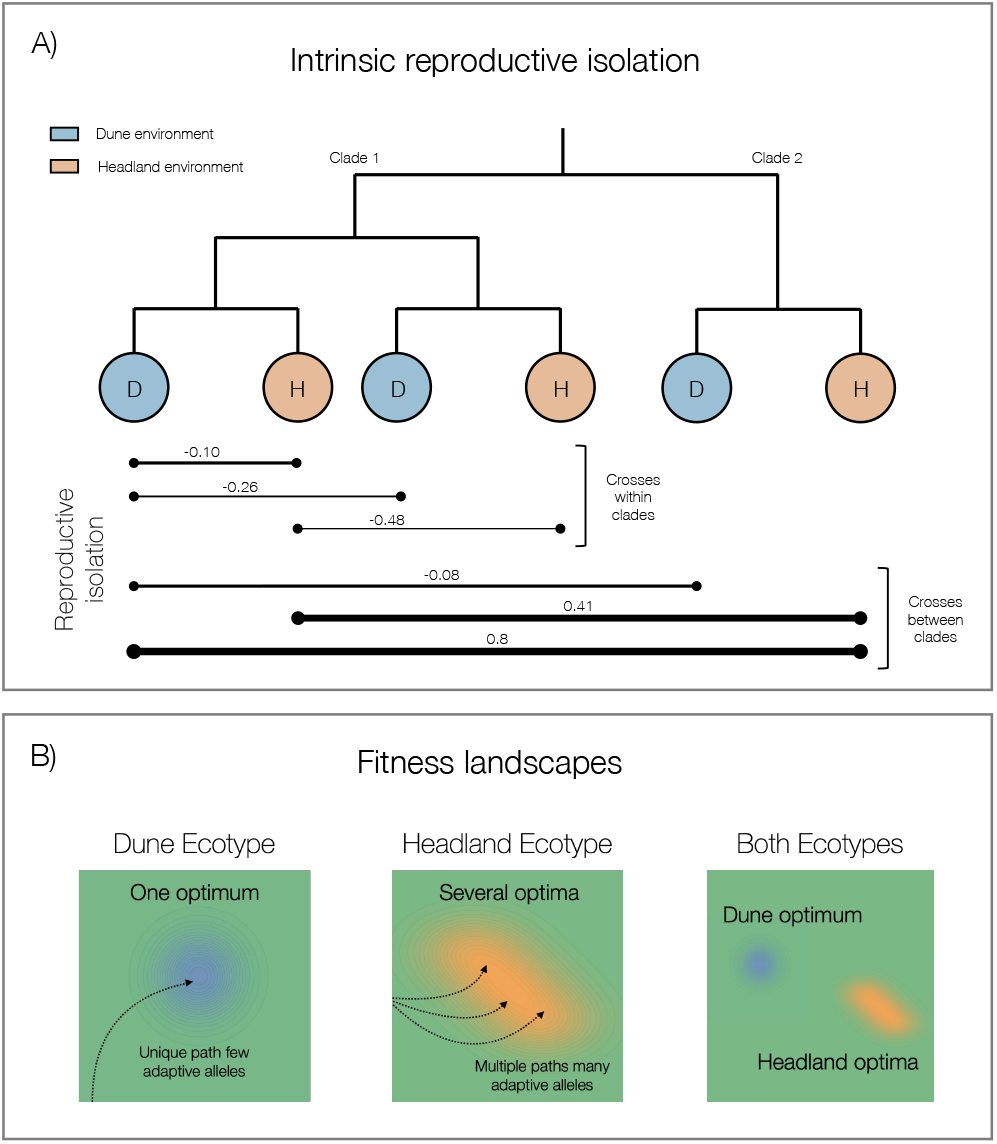
The dynamics of parallel mosaic speciation in *Senecio lautus*. A) Patterns of intrinsic reproductive isolation in *Senecio*. Strong intrinsic reproductive isolation has evolved between ecotypes. In contrast, weaker intrinsic reproductive isolation has evolved between Headland (H) populations but not the Dune (D) populations, as shown by the thickness of the reproductive isolation lines. These patterns of intrinsic isolation are apparent for the most divergent crosses (i.e., crosses between clades). Reproductive isolation values are means for the crosses for F_1_ seed set. B) Schematic diagram representing fitness landscapes of the Dune (blue) and Headland (orange) ecotypes. Dune populations share one optimum, driven by uniform selective pressures, whereas the heterogeneous headland environments likely lead to several optima for the Headland ecotype.

When *φ* ≠ 1, our model predicts that stochastic fixation of different alleles can also lead to DMIs, even under partial environmental overlap. This result parallels the findings of Yamaguchi and Otto (2020), which reveal the speed and direction of environmental change are pivotal in promoting speciation, particularly in cases where populations experience similar but shifting selection pressures. Yamaguchi and Otto (2020) further demonstrate that periods of rapid environmental change increase the likelihood of large-effect mutations becoming fixed, thereby enhancing the probability of DMIs between populations. In the context of parallel evolution, loci controlling convergent traits are predicted to exhibit parallel molecular evolution when selection coefficients are highly asymmetric (*s*_*A*_ ≫ *s*_*B*_) and environmental symmetry is minimal (*φ* →, 1). This pattern is supported by systems such as the parallel loss of armor plates in marine sticklebacks (Jones et al. 2012) and mimicry rings in butterflies (Van Belleghem et al. 2017).

### Complex Genetic Architectures Enhance the Probability of Reproductive Isolation

Our extension of the model to polygenic traits (Equation 3, *P*_*DM*, *P*_) reveals that genetic complexity amplifies the probability of reproductive isolation. *P*_*DM*, *P*,_ captures how combinatorial growth in potential interactions amplifies the effects of selection asymmetry (Figure 5B). Specifically, we show that polygenic adaptation amplifies the effects of selection asymmetry, particularly when multiple loci with varying selection coefficients contribute to adaptation. Our framework explicitly accounts for the combinatorial growth of potential incompatibilities with increasing numbers of loci, explaining why polygenic traits readily accumulate DMIs despite weak selection asymmetry. These results align with Yamaguchi and Otto (2020), who show how pleiotropic mutations, which affect multiple traits, are likely to generate incompatibilities under conditions of rapid adaptation. The *Senecio* system exemplifies these findings: Headland populations achieve phenotypic convergence through distinct combinations of alleles at gravitropism and architectural loci, facilitated by conditional neutrality and asymmetric selection (*φ*< 1; James et al. 2023a; Kaur 2024).

Our polygenic framework also aligns with the work of Schneemann et al. (2020), who explore how the geometry of population divergence in a multi-trait space influences hybrid fitness, distinguishing between intrinsic and extrinsic components of reproductive isolation. Schneemann et al.’s observation that complex trait architectures can either enhance or mitigate hybrid incompatibilities, depending on the distribution of adaptive alleles, closely mirrors our findings of how polygenic architectures and adaptive divergence influence the likelihood and strength of hybrid incompatibilities. Furthermore, their distinction between intrinsic and extrinsic isolation supports our observation that mosaic speciation arises from a dynamic interplay between local adaptation and stochastic divergence. However, as with any proposed mechanistic model, loci contributing to adaptation will not contribute to intrinsic reproductive isolation unless they allow the evolution of DMI systems that require conditional neutrality to create a genetic correlation between adaptation and speciation. See *Supplementary Material 3* for a mechanistic model that connects patterns of reproductive compatibility to parallel evolution in *Senecio*.

Our mathematical framework provides insights into how populations move along the adaptive landscape. When *φ* approaches 1, populations traverse similar adaptive landscapes, minimizing opportunities for incompatibility evolution. As *φ* decreases, increasing landscape complexity creates multiple possible adaptive trajectories, facilitating the accumulation of reproductive barriers even between phenotypically convergent populations, or arriving to different adaptive peaks. These theoretical predictions align with empirical observations in several systems, including Caribbean *Anolis* lizards (Mahler et al. 2013), where fine-scale habitat differences create multiple adaptive solutions despite broad-scale ecological similarities, or in bacteria (Nahum et al. 2015) where environmental heterogeneity can drive populations toward distinct adaptive peaks despite experiencing similar selective pressures.

Together, our mathematical framework demonstrates how uniform, similar and divergent selection can simultaneously shape patterns of reproductive isolation during parallel speciation. The framework unifies ecological and mutation-order mechanisms through a single parameter (*φ*) while revealing how genetic architecture modulates the translation of selection asymmetry into reproductive barriers. These results provide a quantitative foundation for understanding the emergence of biodiversity across heterogeneous landscapes such as in systems like *Senecio*, where environmental variation creates complex mosaics of selection pressures, and therefore of speciation during parallel evolution. Future research should extend the model to include additional complexities, such as recombination dynamics, diploid populations, dominance effects and temporal fluctuations in selection pressures.

### Synthesis and Broader Implications

#### Mechanisms Generating Genetic Incompatibilities During Parallel Speciation

During parallel evolution, reproductive isolation can emerge through several key processes. As shown by our parallel mosaic speciation framework, reproductive incompatibilities can arise via the stochastic fixation of alternative beneficial alleles under similar selective pressures (as with mutation-order speciation), or the deterministic fixation of alternative alleles under divergent selective pressures (as with ecological speciation), or a combination of both (parallel mosaic speciation). These forms of speciation, where reproductive isolation accumulates as a byproduct of natural selection, likely underlie the patterns observed in *Senecio*.

Our theoretical framework demonstrates that genetic incompatibilities may arise through the divergence of pleiotropic developmental pathways that affect both adaptive traits and reproductive compatibility. This mechanism generates several predictions: reproductive barriers should involve genes with dual roles in development and reproduction, when selection coefficients are similar across interacting developmental components, populations can reach similar phenotypic outcomes through different genetic architectures, and hybrids should show a genetic correlation between adaptive and reproductive isolation traits. The *Senecio* system is consistent with this mechanism. Previous work in *Senecio* has found that hormone signaling pathways, particularly auxin regulation, appear to coordinate adaptive traits (gravitropism, architecture; Wilkinson et al. 2021; James et al. 2023a; Broad et al. 2024) and reproductive processes (fertilization success; Wilkinson et al. 2021). Because hormone pathways involve multiple interacting components (McGlothlin and Ketterson 2008) with potentially similar selection coefficients, they provide opportunities for selection to generate different genetic solutions to the same challenges, as seen in other systems of replicated evolution (e.g., Rokas and Carroll 2008; Reid et al. 2016; Birkeland et al. 2020). This integration of deterministic and stochastic processes might explain how *Senecio* populations achieve similar phenotypes through different genetic architectures while accumulating reproductive barriers.

Reproductive isolation can also accumulate between populations via neutral process (True and Haag 2001; Gavrilets 2004; Palmer and Feldman 2009). For instance, system drift is a form of genetic drift where populations traverse a “fitness ridge” of functionally equivalent genetic networks. As populations move along this ridge via stochastic processes, they may develop genetic incompatibilities while maintaining their phenotype and fitness (True and Haag 2001; Schiffman and Ralph 2021; James et al. 2022). Under this process, we expect three key patterns: no systematic relationship between reproductive isolation and ecological divergence, stronger isolation in populations with smaller effective sizes, and random accumulation of incompatibilities with respect to adaptive traits. Our data provide limited support for this mechanism in *Senecio*. Reproductive isolation shows consistent ecotype-dependence rather than random accumulation, effective population sizes remain large across ecotypes (James et al. 2021a), and Headland populations exhibit clear signatures of local adaptation rather than drift. Furthermore, most candidate speciation genes in plants (Rieseberg and Blackman 2010) do not support system drift as a primary driver. Instead, they reflect deterministic evolution due to selection on ecological and reproductive traits. However, specific cases, such as certain S-RNase self-incompatibility genes controlling unilateral incompatibility (Murfett et al. 1996; Hancock et al. 2003) and rare cytoplasmic male sterility loci (Bentolila et al. 2002; Wang et al. 2006), could represent drift-driven incompatibilities, particularly in species or populations with small effective sizes. Even so, these examples are exceptions rather than the rule, suggesting that drift plays a limited role in the evolution of reproductive isolation in plant systems.

Genetic conflict and sexual selection are key mechanisms known to generate intrinsic incompatibilities in plants (Moore and Pannell 2011; Crespi and Nosil 2013; Tonnabel et al. 2021; Coughlan 2023). This can arise through divergent evolution of mating systems (e.g., Martin and Willis 2007), parent-of-origin effects on hybrid fitness (e.g., Coughlan et al. 2020), or antagonistic coevolution between male and female reproductive traits (e.g., Arnqvist et al. 2000). Recent work on conspecific pollen precedence in *Senecio* revealed that local adaptation influences competitive gametic interactions controlled by females (Arenas-Castro 2022), suggesting sexual selection might contribute to reproductive isolation. However, we found no significant asymmetries in hybrid viability or fertility that would indicate strong parent-of-origin effects. The absence of pronounced reciprocal cross differences suggests that while sexual selection may fine-tune reproductive barriers, it is unlikely to be the primary driver of isolation in this system.

#### Release from Trade-offs and the Evolution of Reproductive Isolation

Trade□offs arise when a set of alleles or traits that confer advantages in one environment impose costs in another (Stearns 1989). During the colonization of a new habitat, these constraints can be relaxed or entirely lifted if the selective context shifts, thereby freeing previously suboptimal or deleterious alleles to persist or even become beneficial (Lahti et al. 2009). In the extended Unckless–Orr model, this release is captured mathematically by allowing the parameter *φ* to deviate from 1. When *φ* is significantly less than 1, ancestral constraints no longer fully operate in the new habitat, enabling novel sets of mutations to fix without incurring previous fitness penalties. This release from ancestral trade□offs can accelerate adaptation because it expands the genetic “toolbox” populations have at their disposal. As a result, different populations can fix different beneficial alleles for what appears to be the same ecological task, a phenomenon seen particularly in Headland environments of *Senecio*. Despite convergent prostrate morphologies, Headland populations have accumulated genetic differences that reduce hybrid fitness. These apparently paradoxical patterns arise when the new habitat only partially resembles the ancestral one, making an intermediate value rather than strictly 1 or 0. Such intermediate environmental similarity provides enough shared selective pressures for similar phenotypes to evolve while simultaneously allowing distinct adaptive trajectories, ultimately yielding a mosaic of reproductive barriers across the range. In this way, the relaxation of trade□offs not only promotes phenotypic innovation but can also generate the conditions necessary for the formation of DMIs under parallel mosaic speciation.

#### Effect of Gene Flow on the Likelihood of Parallel Speciation

Population connectivity further complicates the dynamics of speciation because gene flow between populations homogenizes allele frequencies and reduces the likelihood of independent fixation of alternative beneficial alleles (Bank et al. 2012). For example, in populations adapted to similar environments, mutation-order speciation is challenging under moderate gene flow (Schluter 2009; Nosil and Flaxman 2011). This is because advantageous mutations are likely to spread and fix in all populations, thereby preventing the accumulation of incompatibilities. Consequently, mutation-order speciation typically requires very low or absent gene flow (Nosil and Flaxman 2011; Nosil 2012). In *Senecio*, the negligible gene flow in the system (James et al. 2021a) has created ideal conditions for intrinsic incompatibilities to accumulate among Headland populations.

During ecological speciation, where alternative alleles are favored in different environments, there is an antagonism between gene flow and divergent natural selection (Felsenstein 1981). As with mutation-order speciation, gene flow constrains the evolution of reproductive barriers during adaptation, yet speciation remains possible if mechanisms arise to resolve this conflict (also see (Porter and Johnson 2002). If locally adaptive alleles reside in genomic regions of reduced recombination, such as chromosomal inversions, population divergence and speciation can proceed despite ongoing gene flow (Noor et al. 2001; Rieseberg 2001; Butlin 2005; Kirkpatrick and Barton 2006; Ortiz-Barrientos et al. 2016). The role of chromosomal rearrangements in mitigating the antagonism between gene flow and selection has been demonstrated across numerous systems including sticklebacks (Samuk et al. 2017), Atlantic cod (Berg et al. 2017) *Heliconius* butterflies (Martin et al. 2019), *Mimulus* monkeyflowers (Lowry and Willis 2010), *Littorina* snails (Le Moan et al. 2024) and *Drosophila* (Poikela et al. 2024). In the *Senecio* system, the low levels of contemporary gene flow between the Dune and Headland ecotypes likely facilitated the evolution of incompatibilities, yet ancient gene flow may also have shaped their divergence (James et al. 2021a). Future work will explore the role of chromosomal inversions during parallel evolution in *Senecio*, shedding light on their contributions to speciation dynamics.

#### Patterns of Intrinsic Reproductive Isolation across Traits and Development

Reproductive isolation can manifest differently across traits and developmental stages due to varying selective pressures and genetic complexities (Cutter 2023). For instance, while natural selection may act divergently at the population level, individual traits may experience selection that ranges from uniform to highly divergent (Langerhans and Riesch 2013). Additionally, the type and strength of selection upon traits may change over the course of development of the organism, such as when trade-offs exist where traits favoured early in development are not beneficial in later life (Stearns 1989; Garland et al. 2022). Finally, shared genetic architectures among traits can impose pleiotropic constraints, shaping how reproductive barriers evolve and influencing the potential for independent adaptation of specific traits (Mauro and Ghalambor 2020; Zhang 2023).

In *Senecio*, we found that reproductive barriers manifest differently in seed set versus viability. Crosses between Headland populations from different clades show reduced F_1_ seed set, supporting our theoretical predictions about DMI formation under similar selection pressures. However, hybrid viability does not follow this pattern, suggesting that different components of reproductive isolation may evolve through distinct genetic architectures. This asymmetry in reproductive barriers aligns with our mathematical framework: the probability of parallel DMI formation critically depends on both the number of interacting loci and the distribution of selection coefficients. Seed set may be governed by a small number of loci with large effect sizes, such as those involved in specific pollen-pistil recognition and hormone signaling pathways, making parallel selection more likely to generate consistent reproductive barriers. In contrast, hybrid viability likely involves a greater number of loci with smaller, more diffuse effects across genetic networks with an extensive repertoire to disrupt development, increasing the likelihood of different outcomes across the population pairs. This distinction between reproductive barriers requires further exploration to help explain if different components of isolation can evolve through distinct mechanistic pathways even under similar selective pressures.

#### A Continuous Framework for Speciation

Our work expands our understanding of how incompatibilities arise during uniform selection, proposing a broader concept of mutation-order speciation reframed as stochastic speciation. This reframing incorporates some degree of environmental heterogeneity as well as the dynamics of conditional neutrality, emphasizing not only the sequence in which beneficial mutations fix but also the potential for different beneficial alleles to achieve similar adaptive outcomes in comparable environments – an aspect often overlooked in traditional theories of mutation-order speciation. While pure mutation-order speciation can theoretically occur under identical environments, it becomes increasingly improbable with increasing genetic complexity. Most apparent cases of speciation during adaptation to “identical” environments likely involve interactions among multiple mechanisms, including alternative adaptive solutions and subtle environmental heterogeneity. This may explain why the most convincing cases of mutation-order speciation arise in experimental systems (Ono et al. 2017; Hsu et al. 2024), where environmental conditions can be controlled to ensure they are effectively identical. To fully understand the relative contributions of stochastic fixation versus ecological adaptation in natural systems, identifying the specific loci underlying reproductive barriers and their link to local adaptation is crucial (Wu and Ting 2004; Lowry et al. 2008; Nosil and Schluter 2011). Only by characterizing the molecular basis of incompatibilities can we determine whether they arose through the random fixation of alternative beneficial alleles or through adaptation to subtle ecological differences. This will help clarify the extent to which apparent cases of mutation-order speciation in nature truly reflect stochastic processes versus environmental heterogeneity.

The parallel mosaic speciation framework we propose challenges traditional distinctions between ecological and mutation-order speciation (Schluter and Nagel 1995; Ostevik et al. 2012), positing that these modes represent endpoints of a continuous spectrum rather than discrete processes (Langerhans and Riesch 2013). This aligns with recent work showing how speciation dynamics can exhibit “tipping points” where small changes trigger major transitions, suggesting a continuous but potentially nonlinear process (Nosil et al. 2017). Furthermore, local adaptation inevitably involves some degree of reproductive isolation, further supporting this continuous view of speciation (Butlin and Faria 2024). Our framework illustrates how reproductive isolation progresses across gradients of environmental similarity: from deterministic adaptation in identical environments, to adaptation driven by subtle environmental heterogeneity, to divergent adaptation in contrasting environments, where reproductive isolation becomes nearly inevitable. By incorporating environmental symmetry and genetic architecture, our mathematic models help predict speciation outcomes in systems with partial reproductive isolation or varying degrees of environmental heterogeneity. Explicitly treating these factors as continuous variables moves beyond the simple dichotomies of parallel versus divergent evolution towards a more nuanced understanding of speciation mechanisms.

Recent theoretical advances, such as the “speciation hypercube” framework (Bolnick et al. 2023; Johannesson et al. 2024), emphasize the multidimensionality of speciation, integrating genetic, phenotypic, ecological, and temporal factors. In contrast to the traditional “speciation continuum” model (Seehausen et al. 2014; Stankowski and Ravinet 2021), our mathematical analysis complements these multidimensional perspectives by highlighting how parameter-level interactions—including environmental symmetry, selection coefficients, and genetic architecture— generate quantitatively predictable patterns of reproductive isolation. This mechanistic approach bridges descriptive speciation models and evolutionary processes, offering insights into the relative contributions of deterministic versus stochastic processes in driving divergence. We urge researchers to move beyond the categorical classification of ecotypes and instead incorporate within-ecotype environmental variation to better understand the role of environmental heterogeneity in the evolution of reproductive incompatibilities.

## CONCLUSIONS

Our study demonstrates how multiple forms of selection jointly drive the parallel evolution of reproductive isolation in *Senecio lautus*. By integrating field experiments, genetic analyses, and an updated theoretical framework, we reveal that divergent selection between contrasting environments, uniform selection in identical dune environments and similar selection in headland environments together shape speciation dynamics. These findings of parallel mosaic speciation challenge the traditional dichotomy between ecological and mutation-order speciation, highlighting their complementary roles in generating biodiversity.

Incorporating polygenic adaptation and environmental symmetry into our mathematic framework demonstrates that genetic complexity increases the probability of speciation, even without stark ecological contrasts. These findings underscore the importance of genetic architecture in shaping evolutionary outcomes and reveals how phenotypically convergent populations can obscure profound genetic divergence. Future work integrating genomic and ecological data across broader landscapes will further illuminate how divergent and stochastic processes jointly drive the origins of species. Our findings contribute to a growing understanding of how biodiversity emerges and persists across heterogeneous environments.

## AUTHOR CONTRIBUTIONS

Conceptualization: Daniel Ortiz-Barrientos. Formal Analysis: Maddie James, Maria Melo, and Greg Walter. Mathematical Model: Daniel Ortiz-Barrientos and Jan Engelstädter. Experiments: Maria Melo, Maddie James, Federico Roda, Diana Bernal-Franco, Melanie Wilkinson, and Huanle Liu. Data Curation: Maddie James, Maria Melo. Visualization: Maddie James, Greg Walter, and Daniel Ortiz-Barrientos. Writing: Maddie James, and Daniel Ortiz-Barrientos, with contributions from Greg Walter, Melanie Wilkinson, Maria Melo, Federico Roda, Diana Bernal-Franco, Huanle Liu, and Jan Engelstädter. Supervision: Daniel Ortiz-Barrientos. Funding Acquisition: Daniel Ortiz-Barrientos and Jan Engelstädter.

## ACKNOWLEDGEMENTS

The authors acknowledge the Turrbal and Yuggera, Ningy Ningy, Barunggam, Bundjalung, Gumbaynggirr, Gungarri, Gunaikurnai, Gunditjmara, Wirangu, Paredarerme, and Tyerrernotepanner peoples, the traditional owners of the land on which this work was undertaken, and pay their respects to Elders past, present, and emerging. We thank Mohamed Noor, Kieran Samuk, Kate Ostevik, Loren Rieseberg, Eric Baack, Matthew Hahn, and Thomas Richards for their critical discussions on the empirical foundations of this work. We acknowledge the use of ChatGPT v4 Turbo and Claude v3.5 Sonnet assist with minor grammatical edits to the main text, and code development for visualization of the theoretical framework. This research was supported by grants from the Australian Research Council to Daniel Ortiz-Barrientos (DP120104559), Daniel Ortiz-Barrientos and Jan Engelstädter (DP190103039), and Daniel Ortiz-Barrientos through The Australian Research Council Centre of Excellence for Plant Success in Nature and Agriculture (CE200100015).

## CONFLICTS OF INTEREST

The authors declare no conflicts of interest.

## DATA ACCESSIBILITY

All data sets and code will be uploaded to GitHub upon acceptance of the manuscript.

## Notes

### Competing Interest Statement

The authors have declared no competing interest.

### Summary of Updates

After a thorough investigation of the data, both parallel ecological and mutation-order speciation mechanisms appear to be at play in the Senecio system. We show they act together as part of a continuum we call 'parallel mosaic speciation.' We have also added a theoretical framework to understand parallel mosaic speciation.

