## Supplementary Material for "Parallel Mosaic Speciation via Mutation-order and Ecological Divergence"

#### Supplementary Material 1: Methods

To undertake targeted re-sequencing of *Senecio* nuclear genomic regions, we first designed primers in BatchPrimer3 v1.0 (You et al. 2008) using the *S. lautus* contig genome assembly (Liu 2015). We used the default settings with the following adjustments: primer Tm = min 63, opt 64, max 65; primer GC% = 40-80; product size = min 300, opt 400, max 500. We constructed libraries with the Fluidigm® Access ArrayTM system (Moonsamy et al. 2011), which enables the simultaneous amplification of multiple genomic regions across multiple individuals, whilst barcoding each individual with 454 barcodes using the 4-primer PCR process. Pools of amplicons per individual were quantified with the Agilent 2100 Bioanalyzer. Individuals were randomized and combined in equimolar quantities into two pools, and sent for sequencing at the Beijing Genomics Institute. Pools were cleaned with an Agencourt AMPure XP purification kit (removing fragments <150bp), and then sequenced using the emPCR (Lib-A) kit for bi-directional sequencing on two lanes of the Roche GS FLX Titanium platform.

We used the standalone version of TagCleaner v0.12 (Schmieder et al. 2010) to trim forward and reverse barcodes, and remove reads that had > 40% low quality bases (<Q20), > 2% Ns, or were < 50bp. The PRGmatic pipline v1.6 (Hird et al. 2011) was used to align reads into loci and construct haplotypes for each individual, which used the dependent programs: CAP3 (Huang and Madan 1999), BWA v0.7.2 (Li and Durbin 2009), SAMtools v0.1.19 (Li et al. 2009), and VarScan v2.3.6 (Koboldt et al. 2009). We used default parameters and an overlap of 100bp for read alignment (as suggested for 454 data; Hird et al. 2011). We also systematically tested different parameter values to explore their effects on the creation of haplotypes (as undertaken by McCormack et al. 2012; Zellmer et al. 2012), and found that default parameters produced the expected number of sequenced loci (parameters too stringent or relaxed resulted in the creating of too many or few contigs respectively). BLAT v35 (Kent 2002) was used to map haplotypes from PRGmatic to each expected amplicon. For each amplicon, haplotypes were aligned in MUSCLE v3.8 (Edgar 2004). We only considered a locus for further if haplotypes were assigned to at least six individuals per population, excluding S. madagascariensis due to its smaller sample size. Following this criterion, we identified 26 loci.

We next assessed deviations from neutrality for each locus within each population using a combination of the following tests: Hudson-Kreitman-Aguadé (HKA; Hudson et al. 1987), Tajima's D (Tajima 1989), Fu and Li's D*, and Fu and Li's F* (Fu and Li 1993). Analysis was performed in DNAsp v5 (Librado and Rozas 2009). A locus was retained for phylogenetic and population structure analyses if it was considered neutral via the HKA test for all populations (Zhai et al. 2009). In some cases, we were unable to calculate HKA due to the lack of sequencing data in the outgroup (S. madagascariensis). For these loci we assessed neutrality using a combination of the other three tests, and considered a locus neutral if, for all populations, it passed at least two tests. From this we obtained 13 neutral loci across all populations (Table S3).

### Supplementary Material 2: Mathematical Model

The foundational work by Unckless & Orr (2010) (henceforth the UO model) provides a clear approach to mutation-order speciation via Dobzhansky-Muller Incompatibilities (DMIs) under strong selection and weak mutation. Here, we review the UO model and then extend it to more realistic scenarios:

- **Environmental Asymmetry:** Phenotypes may be strongly selected in one habitat but only weakly selected or neutral in another.
- **Polygenic Architectures:** Many loci may contribute to divergence, with the potential for multiple DMIs.

These extensions help to get closer to unifying disparate views of speciation (mutation-order, ecological, polygenic) into a single mathematical framework.

#### Review of the Unckless–Orr (UO) Model

##### Setup of the UO Model

In the simplest form of the UO model, two allopatric populations start with the same ancestral genotype $A_0B_0$. Each population can acquire beneficial mutations at two loci, $A$ and $B$, leading to new alleles $A_1$ or $B_1$. The key assumption is that $A_1B_1$ forms DMI.

The fitness landscape of these genotypes is:

$$\begin{aligned}
A_0B_0 &: 1, \\
A_1B_0 &: 1 + s_A, \\
A_0B_1 &: 1 + s_B, \\
A_1B_1 &: 1 - t,
\end{aligned}$$

where $t > 0$ represents the fitness cost of the incompatibility. Under *strong selection* and *weak mutation*, each beneficial allele sweeps to fixation in one population at a time. DMIs arise if population 1 fixes $A_1$ while population 2 fixes $B_1$, eventually creating the incompatible $A_1B_1$ genotype upon secondary contact.

##### Probability of DMI Formation in the UO Model

Let $P(\text{Pop fixes } A_1)$ be the probability that one population eventually fixes $A_1$ (rather than $B_1$). The main assumption is that beneficial mutations appear one at a time and that the probability of fixation is proportional to the selection coefficient.

If the selection coefficients for $A_1$ and $B_1$ in a given population are $s_A$ and $s_B$, then the expected waiting time for $A_1$ to appear and fix is proportional to $(s_A)^{-1}$, while the corresponding time for $B_1$ is proportional to $(s_B)^{-1}$. Hence the probability that $A_1$ fixes first (and thus excludes $B_1$ from that population) is

$$\frac{s_A}{s_A + s_B}.$$

Similarly, the probability population fixes $B_1$ is $s_B/(s_A + s_B)$.

In two populations (labeled 1 and 2), a DMI arises if population 1 fixes $A_1$ while population 2 fixes $B_1$, or vice versa. Therefore,

$$\begin{aligned} P_{DM,U} &= P(\text{Pop1: } A_1)\ P(\text{Pop2: } B_1)\ +\ P(\text{Pop1: } B_1)\ P(\text{Pop2: } A_1) \\ &= \left(\frac{s_A}{s_A + s_B}\right)\left(\frac{s_B}{s_A + s_B}\right)\ +\ \left(\frac{s_B}{s_A + s_B}\right)\left(\frac{s_A}{s_A + s_B}\right) \\ &= \frac{2 s_A\, s_B}{(s_A + s_B)^2}. \end{aligned}$$

**UO Model: Key Points**

- Two populations, each can fix $A_1$ or $B_1$.
- $A_1 B_1$ is incompatible with fitness $1 - t$.
- Probability of DMI formation is $\frac{2\, s_A\, s_B}{(s_A + s_B)^2}$ when $Ns \gg 1$ and $N\mu \ll 1$.
- Equal selection coefficients ($s_A \approx s_B$) maximize the chance of different alleles fixing in each population.

#### Incorporating Environmental Asymmetry

In nature, environments often differ between populations. A beneficial mutation in one habitat might be only weakly beneficial (or neutral) in another. To capture this, let $\varphi \in [0, 1]$ measure the similarity of selection coefficients across environments. If $\varphi = 1$, the two populations share identical selection pressures; if $\varphi = 0$, an allele beneficial in one environment is effectively neutral in the other.

Suppose:

- $s_A$ is the selection coefficient for $A_1$ in its home environment (population 1).
- $s_B$ is the selection coefficient for $B_1$ in its home environment (population 2).
- In the "foreign" environment, the selection coefficient for $A_1$ is $\varphi\, s_A$, and for $B_1$ is $\varphi\, s_B$.

##### Derivation for $P_{DM,E}$

Under strong-selection, weak-mutation assumptions, each population ultimately fixes one of the two alleles based on the selection coefficient it experiences. Hence:

$$P(\text{Pop1 fixes } A_1) = \frac{s_A}{s_A + \varphi\, s_B}, \quad P(\text{Pop1 fixes } B_1) = \frac{\varphi\, s_B}{s_A + \varphi\, s_B}.$$

$$P(\text{Pop2 fixes } A_1) = \frac{\varphi\, s_A}{\varphi\, s_A + s_B}, \quad P(\text{Pop2 fixes } B_1) = \frac{s_B}{\varphi\, s_A + s_B}.$$

The probability of forming a DMI is then

$$P_{DM,E} = \Big(\frac{s_A}{s_A + \varphi\, s_B}\Big)\Big(\frac{s_B}{\varphi\, s_A + s_B}\Big) \;+\; \Big(\frac{\varphi\, s_B}{s_A + \varphi\, s_B}\Big)\Big(\frac{\varphi\, s_A}{\varphi\, s_A + s_B}\Big)$$

$$= \frac{s_A\, s_B\, \big(1+\varphi^2\big)}{\big(s_A + \varphi\, s_B\big)\,\big(\varphi\, s_A + s_B\big)}.$$

##### Boundary Checks and Example

- $\varphi = 1$**:** Identical environments; recovers the UO formula $\frac{2\,s_A\,s_B}{(s_A+s_B)^2}$.
- $\varphi = 0$**:** Completely dissimilar; $P_{DM,E} = 1$ since each population fixes a different allele.

**Environmental Asymmetry: Key Points**

- Introduces $\varphi \in [0, 1]$ to modulate selection coefficients in alternate environments.
- When $\varphi = 0$, each population is guaranteed to fix distinct alleles ($P_{DM,E} = 1$).
- When $\varphi = 1$, recovers the original UO model.
- Intermediate $\varphi$ shows a smooth transition in $P_{DM,E}$.

#### Polygenic Extension

Adaptation typically involves multiple interacting loci. Here, we extend our model of asymmetric selection to incorporate polygenic adaptation by considering $n$ loci that can form DMIs. This extension shows how genetic architecture interacts with selection asymmetry to influence reproductive isolation.

##### Two-Population, Multi-Locus Model

We consider $n$ loci capable of acquiring beneficial mutations that can generate reproductive incompatibilities. Following the Dobzhansky-Muller model, reproductive isolation arises when populations fix alternative beneficial alleles at interacting loci. Because any derived allele combination produces a hybrid incompatibility, only one derived allele can fix per population. This constraint shapes the probability landscape of DMI formation under polygenic architectures.

For mathematical tractability, we assume $n$ is even and that loci $1, \ldots, n/2$ have selection coefficients $s_{i,1} = s_i$ and $s_{i,2} = \varphi s_i$ in populations 1 and 2 respectively, while loci $n/2+1, \ldots, n$ have $s_{i,1} = \varphi s_i$ and $s_{i,2} = s_i$. This captures the scenario where half the loci are under strong selection in population 1 and weaker selection in population 2, and vice versa.

##### Probability of DMI Formation

Under strong selection and weak mutation, each population will fix exactly one derived allele, with the probability of fixing allele $i$ proportional to its selection coefficient. A DMI forms if the populations fix different alleles. Therefore:

$$P_{DM,P} = 1 - P(\text{populations fix same allele})$$

$$= 1 - \sum_{i=1}^{n} \frac{s_{i,1}}{\sigma_1}\frac{s_{i,2}}{\sigma_2},$$

where

$$\sigma_1 = \sum_{i=1}^{n} s_{i,1}, \quad \sigma_2 = \sum_{i=1}^{n} s_{i,2},$$

are the summed selection coefficients in each population.

For our specific model where half the loci have $(s_i, \varphi s_i)$ and half have $(\varphi s_i, s_i)$, this simplifies to:

$$P_{DM,P} = 1 - \frac{\varphi}{\sigma_1 \sigma_2} \sum_{i=1}^{n} s_i^2$$

**Boundary Cases**

The model yields intuitive results at boundary conditions:

- $\varphi = 0$: Complete environmental divergence yields $P_{DM,P} = 1$, guaranteeing incompatibility.
- $\varphi = 1$: Identical environments recover an $n$-locus version of the UO model:

$$P_{DM,P} = 1 - \frac{\sum_{i=1}^{n} s_i^2}{(\sum_{i=1}^{n} s_i)^2}$$

- Equal selection coefficients ($s_i = s$):

$$P_{DM,P} = 1 - \frac{4\varphi}{n(1+\varphi)^2},$$

  demonstrating that DMI probability increases with locus number and environmental asymmetry.

**Polygenic Extension: Key Points**

- Each population fixes exactly one derived allele to maintain compatibility
- DMI probability depends on both selection coefficient distribution and environmental asymmetry
- Larger numbers of loci increase DMI probability by providing more opportunities for divergence

#### Conclusions

By building on the Unckless–Orr model, we have unified several threads of speciation research:

- **Mutation-order speciation** and **ecological speciation** become endpoints of a continuum governed by $\varphi$.
- **Polygenic architecture** introduces more possible genetic routes to incompatibility, raising the likelihood of speciation.
- **Environmental correlations** shape whether different populations fix the same or distinct alleles, hence controlling whether parallel speciation emerges.

Our results help explain why some species complexes, such as *Senecio lautus* ecotypes, show reproductive isolation patterns consistent with aspects of both parallel mutation-order and ecological speciation.

### Supplementary Material 3: A Mechanistic Model for Parallel Speciation

We develop a mechanistic framework for parallel speciation (Figure and Tables below) that builds upon our extended mathematical model. This framework examines how three loci—**A**, **B**, and **C**—can occupy distinct *functional classes* over evolutionary time, depending on whether they remain in ancestral states or fix derived alleles. In particular:

**1. Universally Adaptive Loci (B and C, ancestral *B, C*).** When $\varphi \approx 1$ (near-identical environments), these loci experience stabilising selection on their ancestral alleles (*B, C*). The probability of forming Dobzhansky–Muller incompatibilities, $P_{\mathrm{DM},E}$, remains low, promoting parallel evolutionary trajectories.

**2. Convergent-Evolution Loci (B and C, derived *b, c*).** Under moderate asymmetry ($0 < \varphi < 1$), the same loci **B**, **C** may fix new, *convergent* alleles (*b, c*) in response to partially shared selective pressures. Despite both populations adopting *b, c*, subtle genetic background differences accumulate, yielding intermediate DMI probabilities. Thus, **B** and **C** appear again in this "second class" because we now focus on their *derived* states.

**3. Conditionally Adaptive Locus (A: *A, a, d*).** This locus can shift between strong or weak selection, depending on the overall genetic context. Even small, per-locus incompatibilities add up across many loci, driving $P_{\mathrm{DM},P}$ higher. When $\varphi < 1$, an initially neutral allele (*a* or *d*) may become strongly selected in a different background, boosting the chance of incompatibilities.

**Why B and C occur in two classes.** They are *the same physical loci*, but their evolutionary role differs based on whether the ancestral (*B, C*) or derived (*b, c*) state is relevant. In near-identical environments ($\varphi \approx 1$), **B** and **C** are "universal" with minimal DMI risk. Under moderate asymmetry ($0 < \varphi < 1$), they fix convergent derived alleles *b, c*, introducing moderate DMIs. Locus **A** is distinct in having multiple conditionally adaptive alleles (*A, a, d*), further multiplying incompatibilities.

Overall, this single set of loci can encompass three *functional* classes of selection and adaptation, rather than requiring separate letters for each class.

**Visual Representation of the Evolution of Hybrid Incompatibilities During Parallel Speciation in *Senecio***

In the figure below, we show how three Dune populations (D1 – D3, blue) independently give rise to three Headland populations (H1 – H3, orange) during adaptation to a new environment. All Dune populations share core adaptive alleles at loci **B** and **C** (*B, C*), which are essential for survival in the ancestral Dune habitat. However, they differ in their genetic backgrounds at locus **A** (*A, a, d*). During adaptation to Headland environments, each population fixes the derived allele *b* at locus **B**, which is associated with the prostrate growth habit, while maintaining their ancestral alleles at loci **A** and **C**.

This divergence creates the potential for DMIs to arise during hybridization due to epistatic interactions between shared adaptive mutations (e.g., *bb*) and divergent genetic contexts at loci **A** and **C**. Specifically:

- **Dune x Dune crosses (D1 x D2, etc.):** These crosses generally result in compatible offspring because they share full ancestral genotypes (e.g., *AABBCC x aaBBCC*). Loci **B** (*BB*) and **C** (*CC*) are core adaptive loci, while locus **A** (*AA or aa*) is neutral within Dunes. Hybrid fitness remains high due to the absence of epistatic incompatibilities.

- **Headland x Headland crosses (H1 x H2, etc.):** These crosses are moderately incompatible due to interactions between fixed derived alleles at locus **B** (*bb*, adaptive) and the divergent genetic backgrounds at loci **A** and **C** (e.g., *AAbbcc x Aabbcc* or *AAbbcc x ddbbcc*). Locus **A** contributes to incompatibilities due to divergence, while locus **C** remains partially neutral. This divergence introduces partial DMIs, reducing hybrid fitness.
- **Dune x Headland crosses (D1 x H1, etc.):** These crosses are highly incompatible because the ancestral alleles at loci **B** and **C** (*BB, CC*, adaptive) in the Dune populations interact negatively with the derived alleles at these loci (*bb, cc*) in the Headland populations. For example, crosses such as *AABBCC x AAbbcc* or *aaBBCC x ddbbcc* exhibit strong genetic incompatibilities. Locus **A** (neutral within ecotypes but divergent between them) exacerbates these incompatibilities, resulting in severe reductions in hybrid fitness.

Arrows in the figure below indicate the evolutionary transitions between populations, with the color coding representing the ecological context (blue for Dunes, orange for Headlands). Letters denote alleles at four interacting loci, where capital letters represent ancestral alleles except for *BB*, which is the ancestral Dune-adapted state, and lowercase letters represent derived alleles except for *bb*, which is the derived Headland-adapted state. The model highlights how reproductive isolation emerges despite phenotypic convergence, primarily due to selection on complex genetic architectures and the interaction of shared and divergent loci during hybridization.

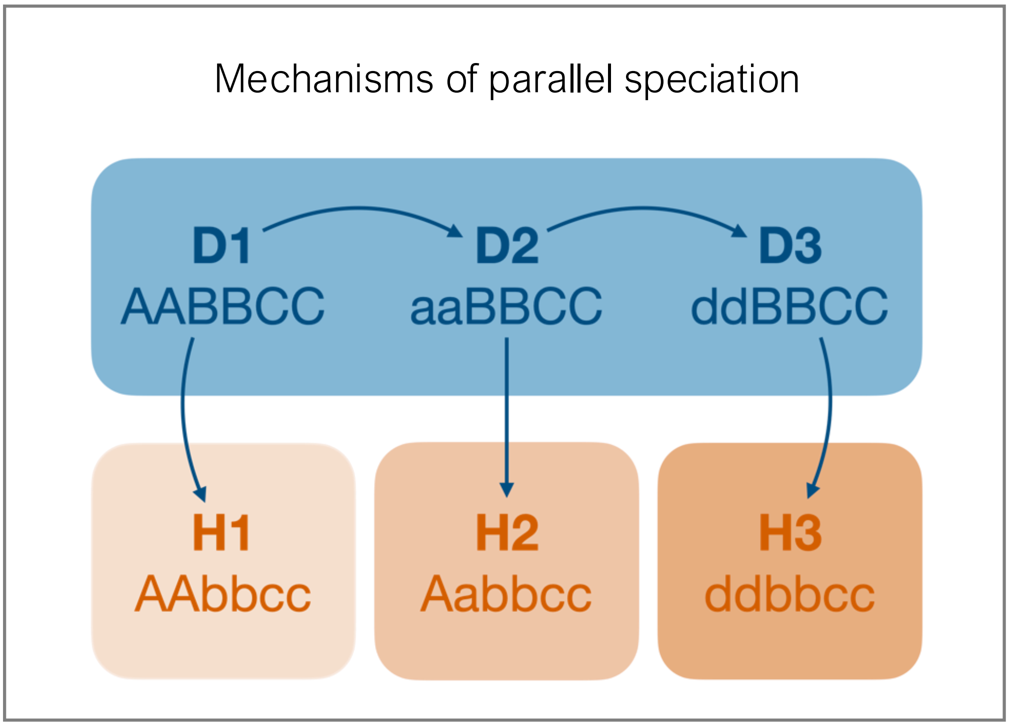

**Crossing relationships between *Senecio* populaitons**

1. **Crosses Between Dunes (DxD):** Dune ecotypes (D1, D2, D3) are compatible and produce high-fitness offspring due to shared genetic backgrounds at most loci. Their fitness remains near 1.0.

2. **Crosses Between Headlands (HxH):** Headland ecotypes (H1, H2, H3) show moderate incompatibilities due to partially divergent genetic architectures at loci A and C. This results in fitness reductions (0.5–0.7), depending on the specific cross.

3. **Crosses Between Dunes and Headlands (DxH):** These crosses produce offspring with very low fitness (0.25–0.4) due to severe incompatibilities between the Dune and Headland genetic backgrounds. Fixation of bb and cc in Headlands interacts negatively with the ancestral Dune alleles (AA, aa, dd).

| | **D1 (AABBCC)** | **D2 (aaBBCC)** | **D3 (ddBBCC)** | **H1 (AAbbcc)** | **H2 (Aabbcc)** | **H3 (ddbbcc)** |
|---|---|---|---|---|---|---|
| **D1 (AABBCC)** | **AABBCC**<br>*Fitness: 1.0* | **AaBBCC**<br>*Fitness: 0.85* | **AdBBCC**<br>*Fitness: 0.85* | **AABbCc**<br>*Fitness: 0.4* | **AABbcc**<br>*Fitness: 0.3* | **AdBbCc**<br>*Fitness: 0.25* |
| **D2 (aaBBCC)** | **AaBBCC**<br>*Fitness: 0.85* | **aaBBCC**<br>*Fitness: 1.0* | **adBBCC**<br>*Fitness: 0.85* | **AaBbCc**<br>*Fitness: 0.4* | **Aabbcc**<br>*Fitness: 0.3* | **adBbCc**<br>*Fitness: 0.25* |
| **D3 (ddBBCC)** | **AdBBCC**<br>*Fitness: 0.85* | **adBBCC**<br>*Fitness: 0.85* | **ddBBCC**<br>*Fitness: 1.0* | **AdBbCc**<br>*Fitness: 0.4* | **Adbbcc**<br>*Fitness: 0.3* | **ddBbCc**<br>*Fitness: 0.25* |
| **H1 (AAbbcc)** | **AABbCc**<br>*Fitness: 0.4* | **AaBbCc**<br>*Fitness: 0.4* | **AdBbCc**<br>*Fitness: 0.4* | **AAbbcc**<br>*Fitness: 1.0* | **AAbbcc**<br>*Fitness: 0.7* | **Adbbcc**<br>*Fitness: 0.5* |
| **H2 (Aabbcc)** | **AABbcc**<br>*Fitness: 0.3* | **Aabbcc**<br>*Fitness: 0.3* | **Adbbcc**<br>*Fitness: 0.3* | **AAbbcc**<br>*Fitness: 0.7* | **Aabbcc**<br>*Fitness: 1.0* | **Adbbcc**<br>*Fitness: 0.5* |
| **H3 (ddbbcc)** | **AdBbCc**<br>*Fitness: 0.25* | **adBbCc**<br>*Fitness: 0.25* | **ddBbCc**<br>*Fitness: 0.25* | **Adbbcc**<br>*Fitness: 0.5* | **Adbbcc**<br>*Fitness: 0.5* | **ddbbcc**<br>*Fitness: 1.0* |

**Mathematical and mechanistic framework for parallel speciation in *Senecio*.** Example of theoretical model of Dobzhansky-Muller incompatibilities (DMIs) connected to empirical patterns observed in the *Senecio lautus* species complex with asymmetric selection effects. The framework can explain how reproductive isolation emerges through the interaction of selection asymmetry ($\varphi$), selection coefficients ($s$), and genetic architecture ($n$). By integrating these parameters, we can predict patterns of reproductive isolation across populations adapting to similar versus contrasting environments. The model demonstrates how both uniform and divergent selection can simultaneously shape speciation, providing a mathematical foundation for understanding parallel mosaic speciation.

| Component | Mechanistic Model | Mathematical Framework | Mechanistic Connection |
|---|---|---|---|
| Dune Compatibility | *BBCC* shared across Dunes; *A/a/d* neutral | High symmetry, $P_{DM,E}$ low | Selection symmetry maintains similar adaptive alleles |
| Headland Incompatibility | Different backgrounds (*AA/aa/dd*); parallel *BB* → *bb* and *CC* → *cc* transitions | Intermediate symmetry, $P_{DM,E}$ moderate | Selection asymmetry accumulates DMIs despite parallel adaptation |
| Dune-Headland Isolation | *BB* → *bb* and *CC* → *cc* transitions | Strong asymmetry, $P_{DM,E}$ high | Strong selection differences drive divergent adaptation |
| Intrinsic Barriers | Epistatic interactions between loci | $P_{DM,P}$ increases with locus number ($n$) | More interacting loci create more opportunities for incompatibilities |
| Extrinsic Barriers | Environment-dependent selection effects | Selection coefficients ($s_A, s_B$) modified by $\varphi$ | Asymmetric selection determines fitness in each environment |
